# A novel *Arabidopsis* phyllosphere resident *Protomyces* sp. and a re-examination of genus *Protomyces* based on genome sequence data

**DOI:** 10.1101/2019.12.13.875955

**Authors:** Kai Wang, Timo Sipilä, Kirk Overmyer

## Abstract

*Protomyces* is a genus of yeast-like fungi that is currently defined as plant pathogens of only the Umbelliferae (Apiaceae) and Compositae (Asteraceae) family plants. Many *Protomyces* species have been proposed; however, there is a lack of molecular data and available specimens for *Protomyces* spp., just over ten species are officially accepted and only six species are preserved and available in public culture collections for examination. Phylogenetic relationships and species boundaries within this genus remain largely controversial. Recently, we isolated *Protomyces* strains from wild *Arabidopsis thaliana* (*Arabidopsis*), a *Brassicaceae* family plant only distantly related to the accepted *Protomyces* hosts. We have previously sequenced the genomes of all the currently public available *Protomyces* species, together with a strain (SC29) we isolated from the *Arabidopsis* phyllosphere. Genome-wide phylogenetic analysis suggests that SC29 occupies a unique phylogenetic position in the genus *Protomyces*. The SC29 genome has low average nucleotide identity values in comparison with other species genomes. As physiological evidence, SC29 has morphological characteristics and carbon assimilation patterns that distinguish it from the other six *Protomyces* species. Analysis with several nuclear gene phylogenetic markers further confirms SC29 as a novel *Protomyces* species and suggests the *act1* gene DNA sequences can be used together with ITS sequences for the rapid identification of *Protomyces* species. In our previous study, SC29 could persist on the *Arabidopsis* phyllosphere in both short term laboratory and overwinter outdoor garden experiments and *Protomyces* spp. (or OTUs) were found in the *Arabidopsis* phyllosphere at multiple sites in both Finland and Germany. We conclude that SC29 is a novel *Protomyces* species isolated from *Arabidopsis* and propose the name of *Protomyces arabidopsidicola* sp. nov. Additionally, the genus *Protomyces* may not be strictly associated with only Compositae or Umbelliferae family host plants, as evidenced by SC29 and *Protomyces* strains previously isolated from plants in other families. The merging of two *Protomyces* species found to have high genomic similarity (*P. inouyei* and *P. lactucaedebilis*) is also proposed.

## INTRODUCTION

*Protomyces* is a genus of plant pathogenic fungi that cause tumour or gall symptoms within flowers, stems, leaves (especially leaf veins), and petioles on host plants exclusively in the Umbelliferae (Apiaceae) and Compositae (Asteraceae) families (Reddy and Kramer 1975; Kurtzman 2011). Members of the genus *Protomyces* have been previously defined based on the morphology of ascospores and vesicles, the host on which they cause disease, and the tissue within the host where they form reproductive structures (Kurtzman 2011; Reddy and Kramer 1975). It has been stated that *Protomyces* species cause disease on “Apiaceae, Compositae, Umbelliferae, and certain other plants” (Kurtzman and Sugiyama 2015; Kurtzman 2011). This claim was supported by citations of Tubaki (Tubaki, 1957) and Reddy and Kramer (Reddy and Kramer 1975). However, a closer examination shows that the genus *Protomyces*, as it is currently defined, only contains species pathogenic on Compositae and Umbelliferae family plants (Table 1).

**Table 1:**
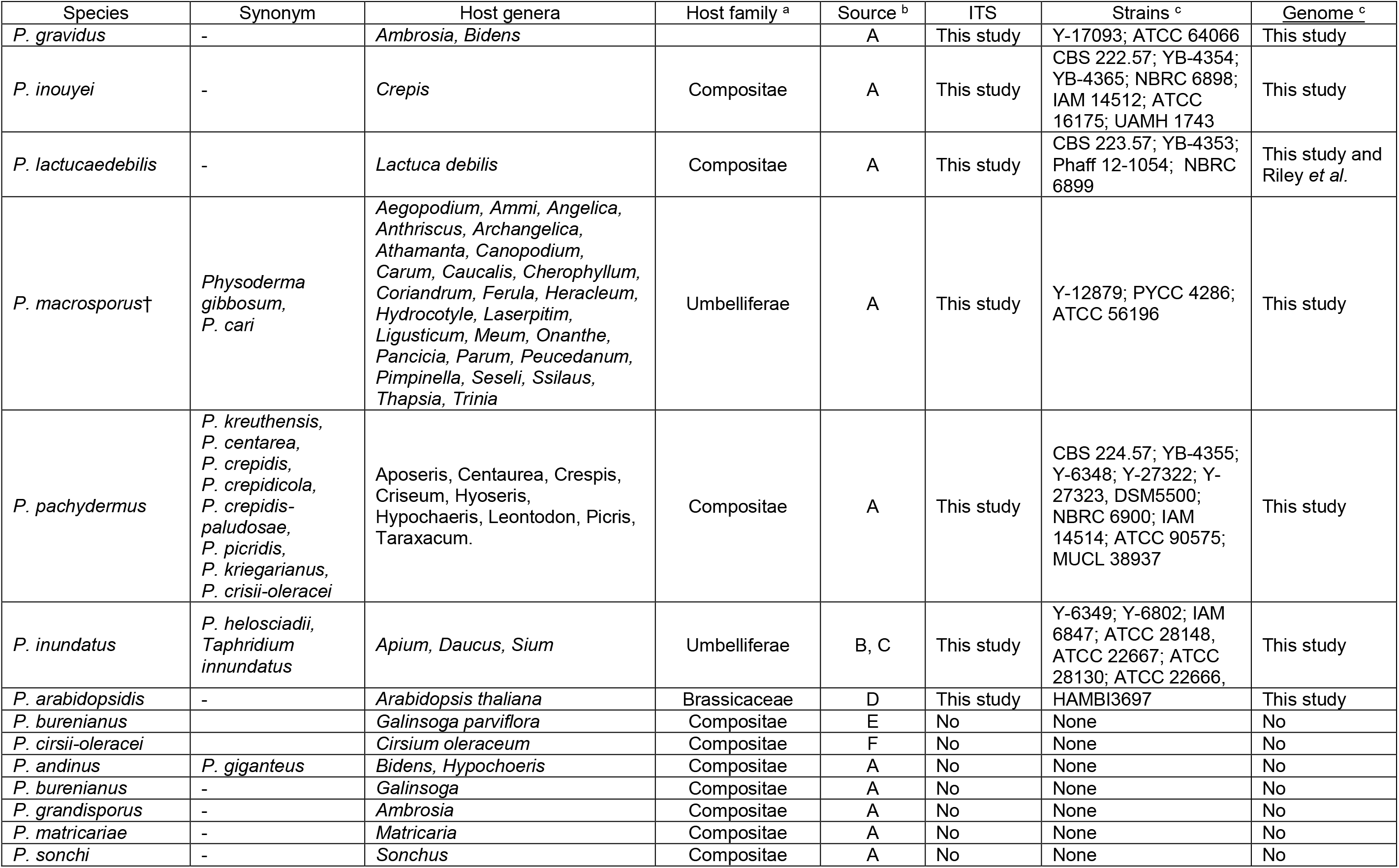
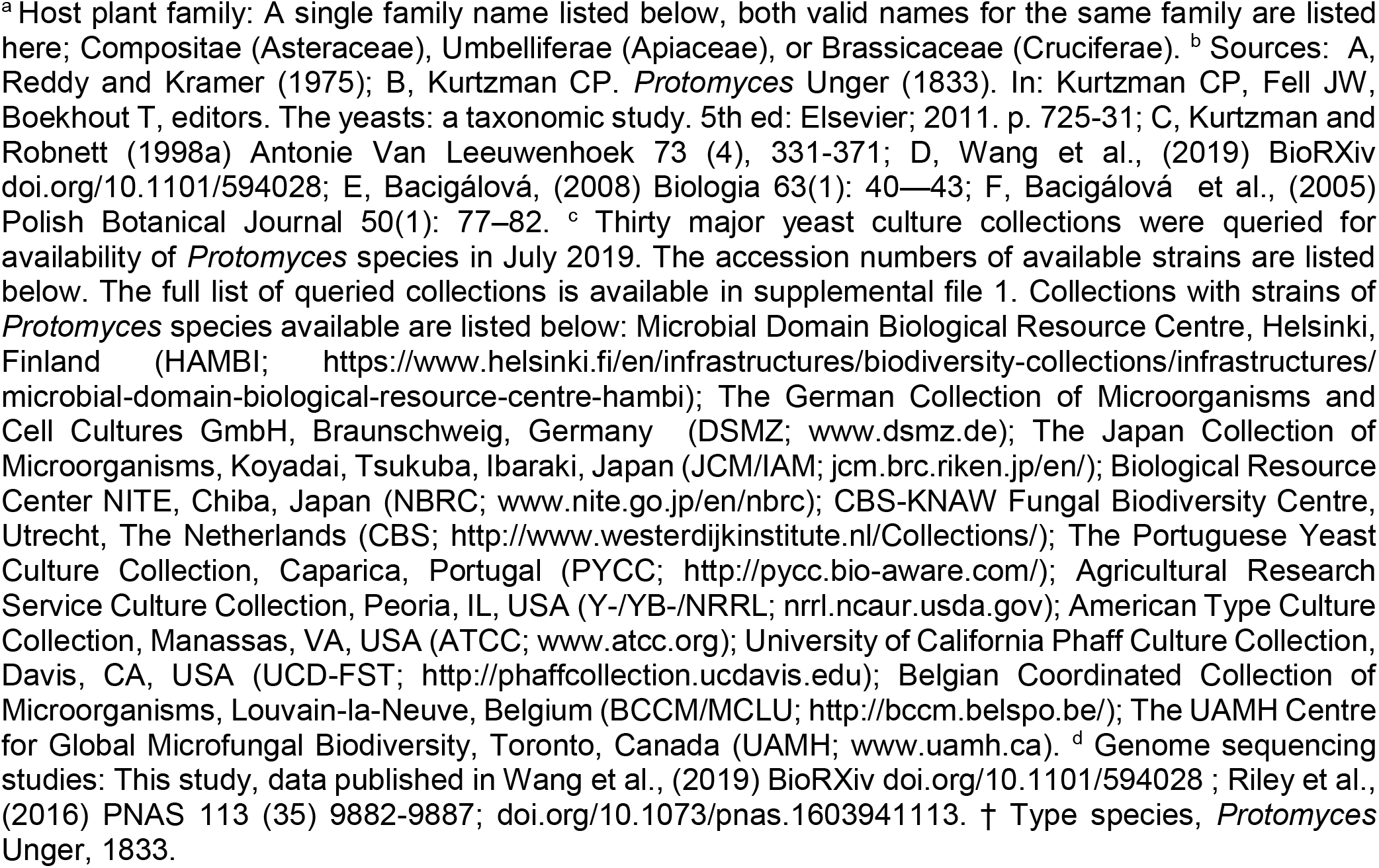
Accepted species of *Protomyces*.

| Species | Synonym | Host genera | Host family <sup>a</sup> | Source <sup>b</sup> | ITS | Strains <sup>c</sup> | Genome <sup>c</sup> |
| --- | --- | --- | --- | --- | --- | --- | --- |
| *P. gravidus* | - | *Ambrosia*, *Bidens* |  | A | This study | Y-17093; ATCC 64066 | This study |
| *P. inouyei* | - | *Crepis* | Compositae | A | This study | CBS 222.57; YB-4354; YB-4365; NBRC 6898; IAM 14512; ATCC 16175; UAMH 1743 | This study |
| *P. lactucaedebilis* | - | *Lactuca debilis* | Compositae | A | This study | CBS 223.57; YB-4353; Phaff 12-1054; NBRC 6899 | This study and Riley *et al.* |
| *P. macrosporus*† | *Physoderma gibbosum*,<br>*P. cari* | *Aegopodium*, *Ammi*, *Angelica*,<br>*Anthriscus*, *Archangelica*,<br>*Athamanta*, *Canopodium*,<br>*Carum*, *Caucalis*, *Cherophyllum*,<br>*Coriandrum*, *Ferula*, *Heracleum*,<br>*Hydrocotyle*, *Laserpitium*,<br>*Ligusticum*, *Meum*, *Onanthe*,<br>*Pancicia*, *Parum*, *Peucedanum*,<br>*Pimpinella*, *Seseli*, *Ssilau*,<br>*Thapsia*, *Trinia* | Umbelliferae | A | This study | Y-12879; PYCC 4286; ATCC 56196 | This study |
| *P. pachydermus* | *P. kreuthensis*,<br>*P. centarea*,<br>*P. crepidis*,<br>*P. crepidicola*,<br>*P. crepidis-paludosae*,<br>*P. picridis*,<br>*P. kriegarianus*,<br>*P. crisii-oleracei* | *Aposeris*, *Centaurea*, *Crepis*,<br>*Criseum*, *Hyoseris*,<br>*Hypochaeris*, *Leontodon*, *Picris*,<br>*Taraxacum*. | Compositae | A | This study | CBS 224.57; YB-4355; Y-6348; Y-27322; Y-27323, DSM5500; NBRC 6900; IAM 14514; ATCC 90575; MUCL 38937 | This study |
| *P. inundatus* | *P. helosciadii*,<br>*Taphridium innundatus* | *Apium*, *Daucus*, *Sium* | Umbelliferae | B, C | This study | Y-6349; Y-6802; IAM 6847; ATCC 28148, ATCC 22667; ATCC 28130; ATCC 22666, | This study |
| *P. arabidopsidis* | - | *Arabidopsis thaliana* | Brassicaceae | D | This study | HAMBI3697 | This study |
| *P. burenianus* | - | *Galinsoga parviflora* | Compositae | E | No | None | No |
| *P. cirsii-oleracei* | - | *Cirsium oleraceum* | Compositae | F | No | None | No |
| *P. andinus* | *P. giganteus* | *Bidens*, *Hypochoeris* | Compositae | A | No | None | No |
| *P. burenianus* | - | *Galinsoga* | Compositae | A | No | None | No |
| *P. grandisporus* | - | *Ambrosia* | Compositae | A | No | None | No |
| *P. matricariae* | - | *Matricaria* | Compositae | A | No | None | No |
| *P. sonchi* | - | *Sonchus* | Compositae | A | No | None | No |

Some confusion may arise from the dual naming of the plant families Apiaceae, which is synonymous with Umbelliferae, and Asteraceae, which is synonymous with Compositae (Turland et al. 2018). The Tubaki paper (Tubaki 1957) is a study of three *Protomyces* species (*P. inouyei, P. lactucaedebilis*, and *P. pachydermus*) pathogenic on Compositae hosts. Reddy and Kramer’s taxonomic revision of the *Protomycetaceae* from 1975 (Reddy and Kramer 1975) clearly indicates that for all the genera in the family *Protomycetaceae*, including all *Protomyces* species, host range is restricted to Umbelliferae and Compositae family plants. This revision work accepted 10 species in the genus *Protomyces*, but rejects 61 previously proposed *Protomyces* species, based on their morphology, lack of materials for examination, or association with host plants outside of the Umbelliferae and Compositae families (Reddy and Kramer 1975). Later works expanded the number of known species in the genus *Protomyces* (Bacigálová 2008; Bacigálová et al. 2005; Kurtzman and Robnett 1998; Kurtzman 2011); however, these species are also all associated with plants within the previously stated restricted host range (Table 1).

There is a clear lack of molecular analysis of species in the *Protomyces* and other *Protomycetales* family genera. Indeed, as has been previously noted (Kurtzman 2011), there is need for increased efforts in collection and molecular analysis of these species. Our survey of 30 major yeast culture collections (Boundy - Mills et al. 2016) revealed that as of July 2019 there are only six *Protomyces* species with strains available for analysis (Table 1).

The genus *Protomyces* was established by Unger in 1833 (Unger 1833) with the type species *Protomyces macrosporus*. The sexual cycle initiates from the diploid hyphal form (Sugiyama et al. 2006), which only occurs during infection in plant tissues. The haploid (asexual) form of *Protomyces* is yeast-like, unicellular, reproduces by budding, and is easy to culture. Carotenoid pigments are formed upon culturing on artificial growth media (Van Eijk and Roeymans 1982) and colonies are usually yellow to pink in colour. The phylogenetic placement of the *Protomyces* has been debated since their first discovery; for review of early work see (Kurtzman 2011). The genus *Protomyces* is the defining member of the family *Protomycetaceae*, which also contains genera with doubtful phylogenetic relationships and genera boundaries (Reddy and Kramer 1975; Kurtzman 2011), including the soil and insect associated *Saitoella*, where the genome of one species has been sequenced (Nishida et al. 2011), and the *Protomyces-like* plant pathogens, *Burenia, Protomycopsis, Taphridium, and Volkartia*, for which there are currently no strains or molecular data available.

Carbon source utilization profiles remain a quick and useful additional tool for the identification yeast species (Kurtzman et al. 2011) and recent fungal genome sequencing projects have begun to address the relationships between biochemical traits and gene content (Riley et al. 2016). However, it was the use of molecular analysis that unambiguously placed the *Protomyces* in the Ascomycete subphylum Taphrinomycotina, as a sister clade to the genus *Taphrina* in the order Taphrinales (Walker 1985; Nishida and Sugiyama 1994; Kurtzman and Robnett 1998; James et al. 2006; Sugiyama et al. 2006; Hibbett et al. 2007; Kurtzman 1993).

It has been noted in the closely related genus *Taphrina* that D1/D2 and especially ITS markers gave resolution to the genus level, but some species could not be resolved (Rodrigues and Fonseca 2003). In general, there is a need in some fungal taxa to develop lineage specific secondary phylogenetic markers to achieve reliable species level identification; DNA markers, such as actin, translation elongation factors, ribosomal polymerase subunits and tubulin, have been described and applied for this purpose (Stielow et al. 2015).

It has been suggested that members of the genus *Protomyces*, its sister genus *Taphrina*, and some other members of the order Taphrinales, may have retained the lifestyles of early Ascomycetes, due their many ancestral features and basidiomycete-like traits, such as high genomic GC content (Sugiyama et al. 1996; Wang et al. 2019), thick walled “chlamydospore” reproductive or resting cells (Bary and Garnsey 1887; Mix 1924), basidiospore-like naked asci (Sadebeck 1884; Lohwag 1934), enteroblastic budding pattern (Sugiyama et al. 1996; Von Arx and Weijman 1979) Q-10 ubiquinone system (Sugiyama et al. 2006), and the presence of a dual hybrid histidine kinase (Wang et al. 2019). These similarities are also illustrated by the many instances, in which species within the order Taphrinales have been misclassified among the Basidiomycetes, or vice versa (Piepenbring and Bauer 1997; Bary and Garnsey 1887; Reddy and Kramer 1975; Nishida and Sugiyama 1995). Due to their phylogenetic position and these characteristics, these species are of considerable evolutionary interest.

Here we describe *Protomyces* sp. strain C29 (SC29), isolated from the phyllosphere of wild *Arabidopsis thaliana*, as a new *Protomyces* species, and propose the name *Protomyces arabidopsidicola* sp. nov. The definition of the genus *Protomyces* and boundaries species within are also examined in light of new genome sequencing data.

## MATERIALS AND METHODS

Six *Protomyces* reference species (Table 1) were from the USDA ARS culture collection (NRRL; https://nrrl.ncaur.usda.gov/). Species/strains used and their abbreviations are: Para, *Protomyces arabidopsidicola* sp. nov.; SC29, Para strain C29; Pgra, *P. gravidus*; Pino, *P. inouyei*; Pinu, *P. inundatus*; Plac, *P. lactucaedebilis*; Pmac, *P. macrosporus*; Ppac, *P. pachydermus*. The isolation and culture *Protomyces* SC29 from the leaf surface of healthy wild-growing *Arabidopsis thaliana (Arabidopsis)*, in Helsinki, Finland, were described in our previous study, Wang *et al*. (Wang et al. 2016). DNA extraction, PCR amplification or marker sequences, and DNA sequencing, were described in our previous studies (Wang et al. 2019; Wang et al. 2016). Average nucleotide identity (ANI) and average amino-acid identity (AAI) values of *Protomyces* genomes or proteomes were calculated using online tool ANI/AAI-Matrix (Rodriguez-R and Konstantinidis 2016). Growth assays at low temperature were previously reported in (Wang et al. 2019). Yeast cell size and morphological characterization was conducted by photographing three-day-old or two-week-old cultures with a LEICA 2500 microscope (www.leica-microsystems.com) or two-week-old colonies with a LEICA MZ10F stereo microscope. Cells and colonies were cultivated on GYP (glucose yeast-extract peptone) agar plates. For cell size measurements yeast cells were mounted on a slide in water for examination. Cell length and width were measured from photomicrographs with imageJ software (https://imagej.nih.gov/ij/). Carbon assimilation patterns among seven species were tested with API 50 CH strips (bioMerieux SA; www.biomerieux.com) cultured at 21 °C for seven days, according to the manufacturer’s instructions.

The genome sequences and annotations of *Saitoella complicata* Saico1, *Saccharomyces cerevisiae* GCF000146045.2 R64, *Schizosaccharomyces pombe* GCF000002945.1, *Pneumocystis murina* B123, *Taphrina populina* JCM22190, *Neolecta irregularis* DAH-3 were downloaded from NCBI (www.ncbi.nlm.nih.gov/). The phylogenetic tree with species representing Taphrinomycotina classes Neolectomycetes, Pneumocystidomycetes, Schizosaccharomycetes, and Taphrinomycetes was built with 636 single-copy protein sequences from genome annotations. *Saccharomyces cerevisiae*, representing Saccharomycotina, was utilized as an outgroup. Conserved single-copy protein sequences were identified with orthofinder (Emms and Kelly 2015). Alignment quality was controlled by applying sequence scores >= 0.8 in MAFFT analysis with Guidance2. Multiple sequences were concatenated with FASconCAT_V1.0. The randomized axelerated maximum likelihood method RAXML (Stamatakis 2014) and rapid bootstrapping (100x) (Stamatakis 2014) were employed for building the genome-wide tree, and (1000x) for nuclear DNA marker trees. Phylogenetic trees using the nuclear DNA markers; actin1 (*act1*), second largest subunit of RNA polymerase II (*rbp2*), large subunit of RNA polymerase II (*rbp1*), transcription elongation factor 1 (*tef1*) and tubulin beta chain (*tub2*) were aligned by ClustalX2.1 (Larkin et al. 2007). Sequences of DNA marker gene orthologs in *Protomyces* and *Taphrina* were discovered using local BLAST searches of their genomes using *Schizosaccharomyces* sequences as query. All phylogenetic trees were viewed and edited with iTOL (Letunic and Bork 2016). Trees based on the Bayesian inference method were constructed utilizing the MrBayes software package (Ronquist et al. 2012), selecting the general time reversible (GTR) model and invgamma. Two independent analyses were started simultaneously with 10000 generations and four chains set for the analysis run.

The sequences of proteins known to be involved in carbohydrate utilization in model yeast species were collected from NCBI and UniProt (https://www.uniprot.org/). Characterized protein sequences of key enzymes were selected as query sequences. Sequences used as search queries are listed in supplemental file 1. A local installation of BLASTp was applied for searching enzyme hits in *Protomyces* genomes. Selection criteria are E value < 1e^−30^ and protein hits were manually curated to avoid duplicates. All analysis above were performed using CSC (https://www.csc.fi/) computing resources in Linux.

## RESULTS

### Evidence of SC29 as a new Protomyces species

We have previously isolated a novel *Protomyces* sp. strain C29 (SC29) from the phylloplane of healthy individuals of thale cress (*Arabidopsis thaliana*; hereafter referred to as *Arabidopsis*) (Wang et al. 2016) growing in the wild in Helsinki, Finland. SC29 (as a member of OTU1) was identified as belonging to the genus *Protomyces* by BLAST searches with ITS sequences (Genbank acc. LT602858) showing highest similarity (96 %) to *P. inouyei* (Wang et al. 2016). The SC29 genome and the genomes of six reference species were previously sequenced and subjected to comparative analysis (Wang et al. 2019). These six sequenced species represent all of the *Protomyces* species, for which strains are currently available in yeast culture collections (Table 1). All sequenced *Protomyces* spp. have small genome sizes (11.5-14.1 Mbp) with 50.9-52.8 % GC content. Our previous work (Wang et al. 2019) focused on the basic description of SC29, its interactions with *Arabidopsis*, and comparison of the features of the seven sequenced genomes. In the current work, we focus on the description and naming of SC29 as a novel species as well as the phylogenetic implications of our *Protomyces* genome sequencing results (Wang et al. 2019).

In Fig. 1, the typical peat moss and stone environmental conditions of sites where *Arabidopsis* samples were collected for yeast isolation is shown (Fig. 1a) as well as a representative individual of the healthy *Arabidopsis* plants collected in their native growth habitat (Fig. 1b). The SC29 haploid yeast stage in culture is presented in Fig. 1c. Yeast cell sizes (Table 2) as well as cell and colony morphology (Fig. 2) of SC29 and all six reference *Protomyces* spp. were analysed, where SC29 cell size was significantly shorter than its closest relative *P. inouyei*, further suggesting it is a distinct species.

**Fig. 1.**
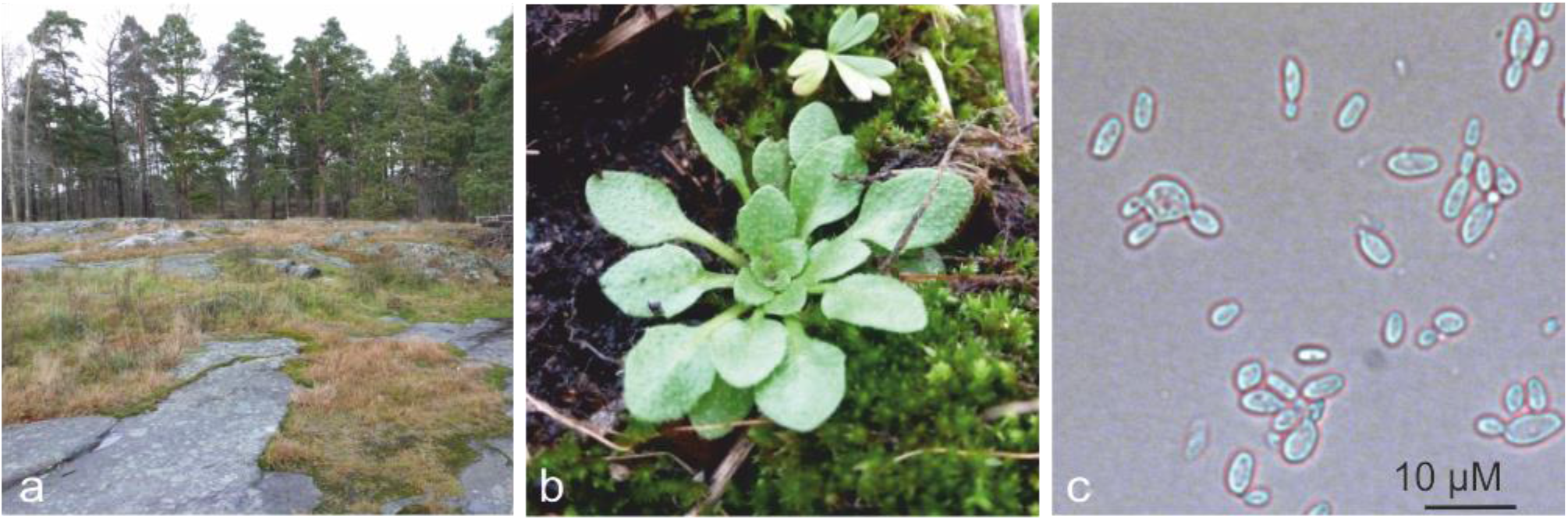
Isolation environment and host. a), The environment from which *Protomyces arabidopsidicola* strain C29 (SC29) was isolated. b), A typical example of healthy wild thale cress (*Arabidopsis thaliana*) plants that were collected for yeast isolation. c), Yeast cell morphology of three-day-old SC29 when cultured on PYG (peptone-yeast extract-glucose) agar plate.

**Fig. 2.**
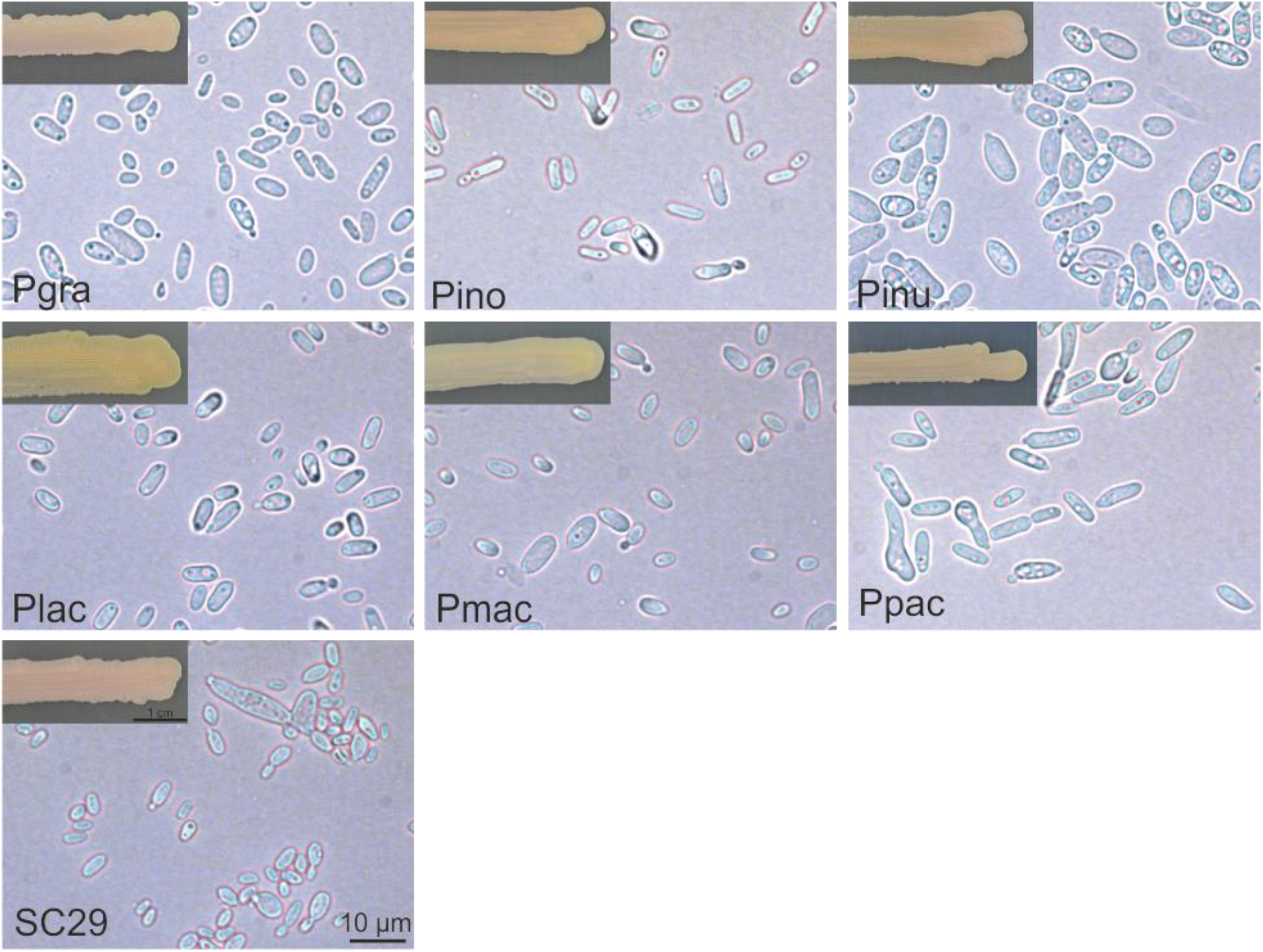
Morphology of *Protomyces spp*. Morphology of yeast cells and colonies of *Protomyces* species including SC29. Yeasts were cultured in GYP (glucose yeast-extract peptone) agar (2 %) medium. Photos of colonies taken two weeks after inoculation. Photos of cells were taken three days after inoculation. Pgra: *P. gravidus*, Pino: *P. inouyei*, Pinu: *P. inundatus*, Plac: *P. lactucaedebilis*, Pmac: *P. macrosporus*, Ppac: *P. pachydermus*, SC29: *P*. sp. SC29. Scale bars in SC29 panel, 10 μm for east cells and 1 cm for colonies, are valid for all panels.

**Table 2.** Yeast cell sizes. Measurements were from three-day- (3 d) and 14-day-old (14 d) cells of *Protomyces* species. Yeast cells were cultured in GYP agar plates. Cell images were captured by LEICA 2500 microscope with camera LEICA DFC490. Cell length and width were measured with imageJ software. Two independent cultures were applied in the statistics, with around 100 cells sampled in each biological repeat. The yeast cells of *P*. sp. SC29 had a significantly shorter length than its closest relative *P. inouyei* (p<0.05) by one way ANOVA + Tukey HSD. NA: not available. “Reference*”: is the published size range for each species according to Kurtzman, 2011.

| Age | Species | Mean ± SD ($\mu\text{m}$) | Size range | Reference* |
| --- | --- | --- | --- | --- |
| 3 d | *P. sp. SC29* | 4.8±1.6 x 2.7±0.6 | 2.0-12.4 x 1.4-5.6 | NA |
| | *P. gravidus* | 5.2±1.3 x 3.0±0.6 | 2.7-9.6 x 1.4-5.9 | 2.5-10 x 2.5-4 |
| | *P. inouyei* | 6.3±1.6 x 2.6±0.4 | 3.2-11.4 x 1.5-4.2 | 2.5-10 x 2-4 |
| | *P. inundates* | 7.5±2.1 x 3.8±0.6 | 3.8-14.2 x 1.1-5.4 | 3.7-9 x 2-4.7 |
| | *P. lactucaedebilis* | 5.4±1.5 x 2.7±0.4 | 3.0-10.8 x 1.6-4.0 | 3.5-9 x 2.5-5 |
| | *P. macrosporus* | 5.2±1.6 x 2.9±0.5 | 2.3-13.0 x 1.7-4.2 | 3-7 x 2.5-4 |
| | *P. pachydermus* | 7.4±2.1 x 2.9±0.6 | 3.6-13.6 x 1.6-5.5 | 3-8 x 2.5-4 |
| 14 d | *P. sp. SC29* | 4.7±1.8 x 2.6±0.8 | 2.4-14.1 x 1.6-5.7 | NA |
| | *P. gravidus* | 5.4±1.4 x 2.6±0.5 | 3.2-8.2 x 1.4-3.8 | |
| | *P. inouyei* | 4.7±1.3 x 2.7±1.1 | 2.6-10.8 x 1.1-6.5 | |
| | *P. inundates* | 8.7±3.8 x 5.6±2.7 | 3.6-18.7 x 1.9-16.2 | |
| | *P. lactucaedebilis* | 5.2±1.5 x 2.9±0.8 | 2.8-9.7 x 1.5-5.0 | |
| | *P. macrosporus* | 4.9±1.3 x 2.7±1.0 | 3.0-8.5 x 1.6-8.1 | |
| | *P. pachydermus* | 6.4±2.6 x 3.0±0.5 | 3.8-20.8 x 1.9-4.0 | |

In order to reveal the evolutionary relationships and species delimitation within the genus *Protomyces*, we built phylogenetic trees using various markers; specifically, DNA sequences of the 26S large-subunit (LSU) ribosomal DNA (rDNA) D1/D2 domain, the full ITS DNA sequences (ITS1-5S rDNA-ITS2), and conserved protein sequences from the genome sequencing data of SC29 and the reference species. All phylogenetic trees were constructed both using the maximum-likelihood method and the Bayesian inference method (Fig. 3), which both yielded similar results and support the same conclusions. The ITS marker differentiated all strains at the species level, but with low support for some nodes (Fig. 3, a and b). D1/D2 trees offered poor species resolution and were also poorly supported (Fig. 3, c and d). These results suggest that the true diversity of the genus *Protomyces* is not captured or supported in the phylogenies utilizing the commonly used D1/D2 or ITS markers. Therefore, other methods are required to resolve the phylogeny of species in the genus *Protomyces*. To this end, we utilized genome sequencing data and performed genome-wide phylogenetic analysis using 1670 single-copy protein sequences that were common to these seven genomes. Clearly, SC29 occupies a unique position in a monophyletic clade with the six *Protomyces* spp., indicating it is a unique species of *Protomyces* most closely related to a clade composed of Pino, Plac, and Ppac (Fig. 3, e and f). As genomic sequencing is not practical for rapid species identification, we tested five nuclear genes (*rbp1*, RNA polymerase subunit 2; *tef1*, translation elongation factor 1; *rbp1*, RNA polymerase subunit 1; *tub2*, tubulin beta; and *act1*, actin 1) as potential secondary lineage specific phylogenetic DNA markers, both individually and together as a single concatenated sequence (Fig. 4). All of these nuclear DNA markers resolved SC29 as distinct from other *Protomyces* species, providing additional evidence at the DNA level that SC29 represents a novel species of *Protomyces* (Fig. 4). All markers performed reasonably well at resolving species in the genus *Protomyces* (Fig. 4).

**Fig. 3.**
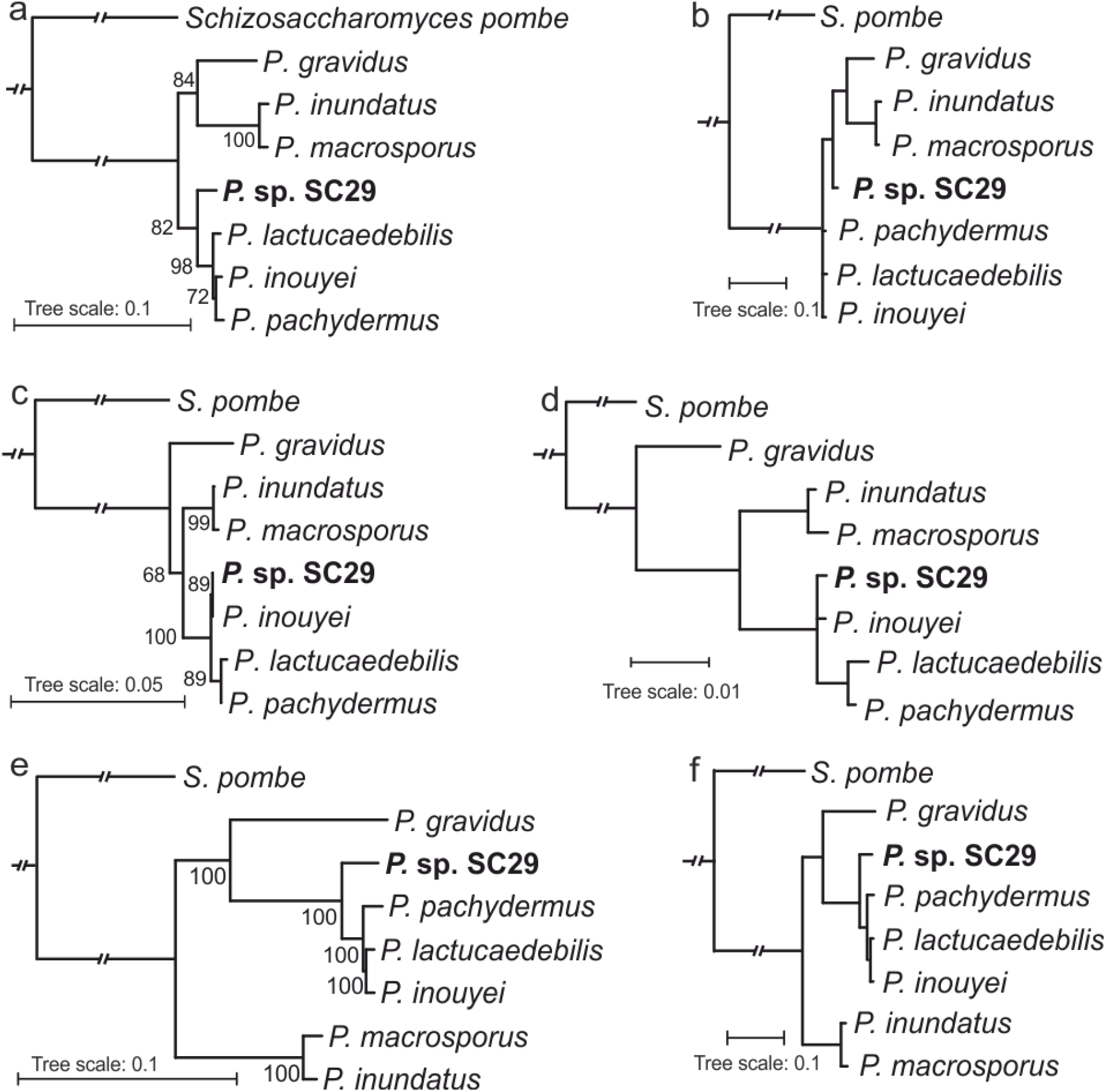
Phylogenetic analysis of the genus *Protomyces*. Phylogenetic trees built by ITS (a, b), D1D2 (c, d) and genome-wide sequences (e, f), with maximum likelihood (a, c, e) and Bayesian method (b, d, f). For maximum likelihood method, a), ITS and c), D1D2 sequences were aligned with ClustalX2. Bootstrapping (1000x) was conducted and best-scoring ML tree were produced with randomized axelerated maximum likelihood (RAxML) rapid bootstrapping program. Bootstrap support values (%) are indicated at each node. e), RAxML with rapid bootstrapping (100x) were chosen for constructing the maximum likelihood tree utilizing 1670 concatenated single-copy conserved protein sequences from whole genome sequence data. Alignment quality control of single-copy conserved proteins was achieved by applying sequence scores >= 0.8 in MAFFT analysis using Guidance2. Multiple aligned sequences of each species/strain were concatenated using FASconCAT_V1.0 and bootstrap values (%) are indicated at the nodes. (b, d, f) Bayesian phylogenetic trees were produced by using program MrBayes version 3.2.7a. The input .nex files are generated from the aligned fasta files used in (a, c, e). General time reversible (GTR) model and invgamma were chosen. Two independent analyses were started simultaneously, and 10000 generations and four chains were set for analysis run. The output .tre files were viewed with online tool iTOL. In all phylogenies *Schizosaccharomyces pombe* was used as an outgroup.

**Fig. 4.**
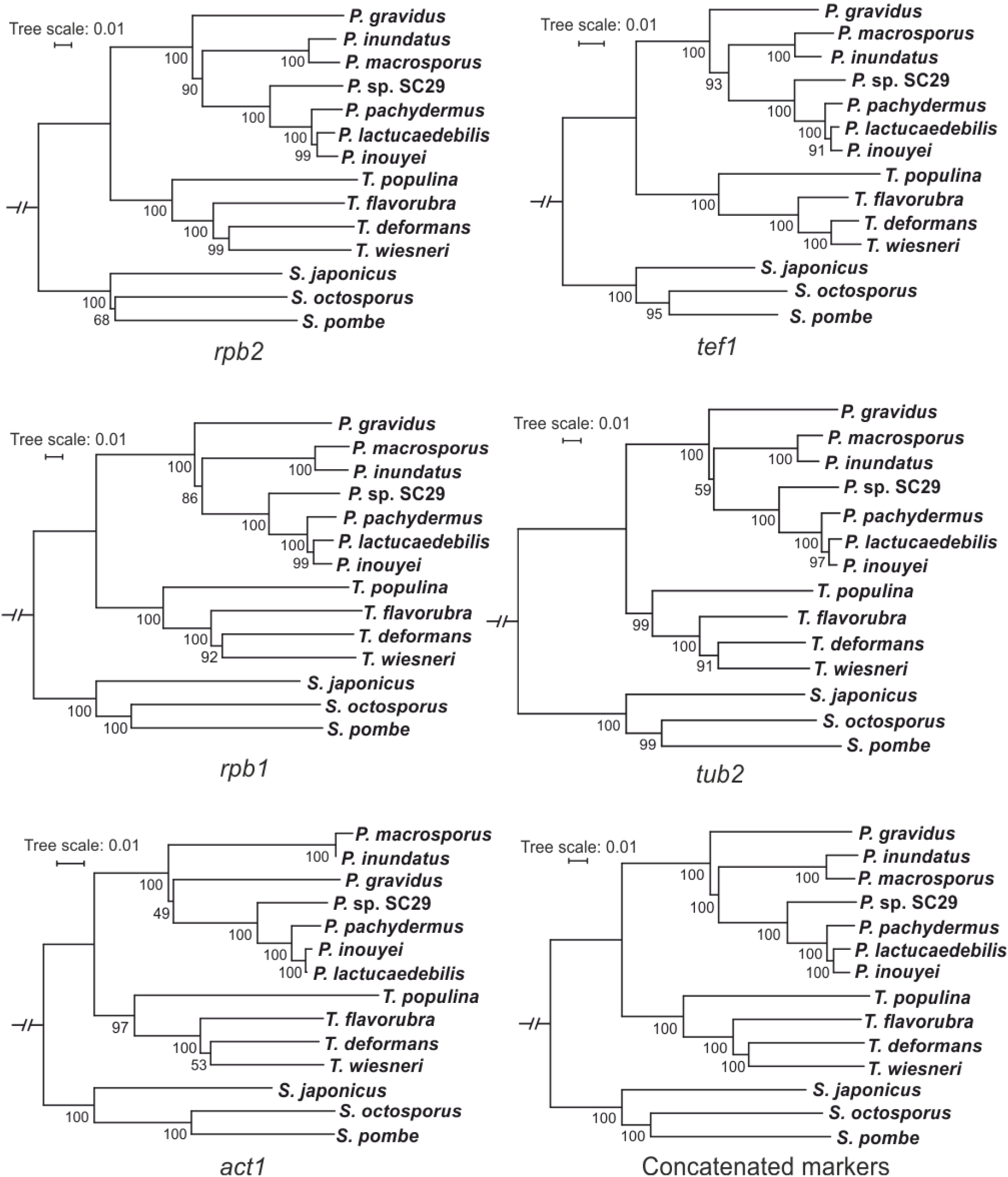
Nuclear marker gene phylogenetic trees. Phylogenetic trees of species in the genera *Protomyces* and *Taphrina* with the nuclear gene markers *rbp2* (RNA polymerase subunit 2), *tef1* (translation elongation factor 1), *rbp1* (RNA polymerase subunit 1), *tub2* (tubulin beta), *act1* (actin 1) and concatenated sequence of the five nuclear marker genes above. Protein sequences from the respective marker genes in *Schizosaccharomyces pombe* were used as a query for BLASTp searches against protein sequences from *Protomyces* and *Taphrina* genome annotations. DNA sequences of each marker were then collected and aligned with ClustalX2 to construct neighbour-joining trees with 1000 bootstraps. Bootstrap support values (%) are indicated at each node. *Schizosaccharomyces pombe* was used as an outgroup.

In addition to phylogenetic analysis, comparisons of whole genome average nucleotide identity (ANI) and average amino acid identity (AAI) between SC29 and the other six species were used as evidence to define species borders. ANI and AAI values were between 77.5 to 85.6 % and 71.9 to 90.6 %, respectively (Table 3). The low levels of ANI and AAI provide further evidence that SC29 is a new species that is distinct from the other six *Protomyces* species. Further comparative genomic analysis of these *Protomyces* spp. are presented elsewhere (Wang et al. 2019).

**Table 3.** ANI and AAI values of *Protomyces* spp. genomes. ANI (average nucleotide identity) and AAI (average amino-acid identity) values among *Protomyces* species. Genome and annotation matrix were applied with online tool ANI/AAI-Matrix using data from (Wang et al. 2019). The lower matrix shows the ANI values (in bold) and the upper matrix indicates the AAI values (in plain text). Species name abbreviations used: *Protomyces inundatus* (Pinu), *P. macrosporus* (Pmac), *P. gravidus* (Pgra), *P. lactucaedebilis* (Plac), *P. inouyei* (Pino), *P. pachydermus* (Ppac), and *P. arabidopsidicola* strain C29 (Para).

|  | Pinu | Pmac | Pgra | Plac | Pino | Ppac | Para |
| --- | --- | --- | --- | --- | --- | --- | --- |
| Pinu | - | 94.7 | 71.4 | 71.9 | 71.8 | 71.9 | 72.0 |
| Pmac | **92.0** | - | 71.3 | 71.8 | 71.7 | 71.9 | 71.9 |
| Pgra | **77.9** | **77.8** | - | 74.2 | 74.1 | 74.1 | 74.2 |
| Plac | **78.0** | **78.3** | **77.6** | - | 97.3 | 95.6 | 90.6 |
| Pino | **78.0** | **77.9** | **77.6** | **96.0** | - | 95.6 | 90.6 |
| Ppac | **78.0** | **77.8** | **77.5** | **93.6** | **93.6** | - | 90.1 |
| Para | **77.9** | **78.1** | **77.5** | **85.6** | **85.5** | **85.0** | - |

Carbon assimilation profiles of SC29 and the other six *Protomyces* species SC29 were determined (Table 4). This data demonstrates that each species utilized a unique pattern of carbon sources. SC29 exhibited a distinct profile of carbon utilization traits, especially for D-cellobiose, amygdalin, L-arabinose and D-arabinose (Table 4) further distinguishing it from *P. inouyei*. Based on this data, a new diagnostic examination of seven species/strains is provided (Table 5). Utilizing the genome annotations of seven *Protomyces* species, we compared the presence of genes known to be involved in carbon source metabolism with selected metabolic traits (Fig. 5), including those that distinguish the *Protomyces* species in this study (Table 4). For some traits such as utilization of D-cellobiose, L-rhamnose, D-ribose, and D-mannitol, the presence or absence of full pathways and the number of paralogs correlated with metabolic phenotypes (Fig. 5). However, for amygdalin, L-arabinose, D-arabinose, and D-xylose, the genomically encoded enzymes were generally highly conserved in all species and thus do not correlate well with the inability of some species to utilize these carbon sources (Fig. 5).

**Fig. 5.**
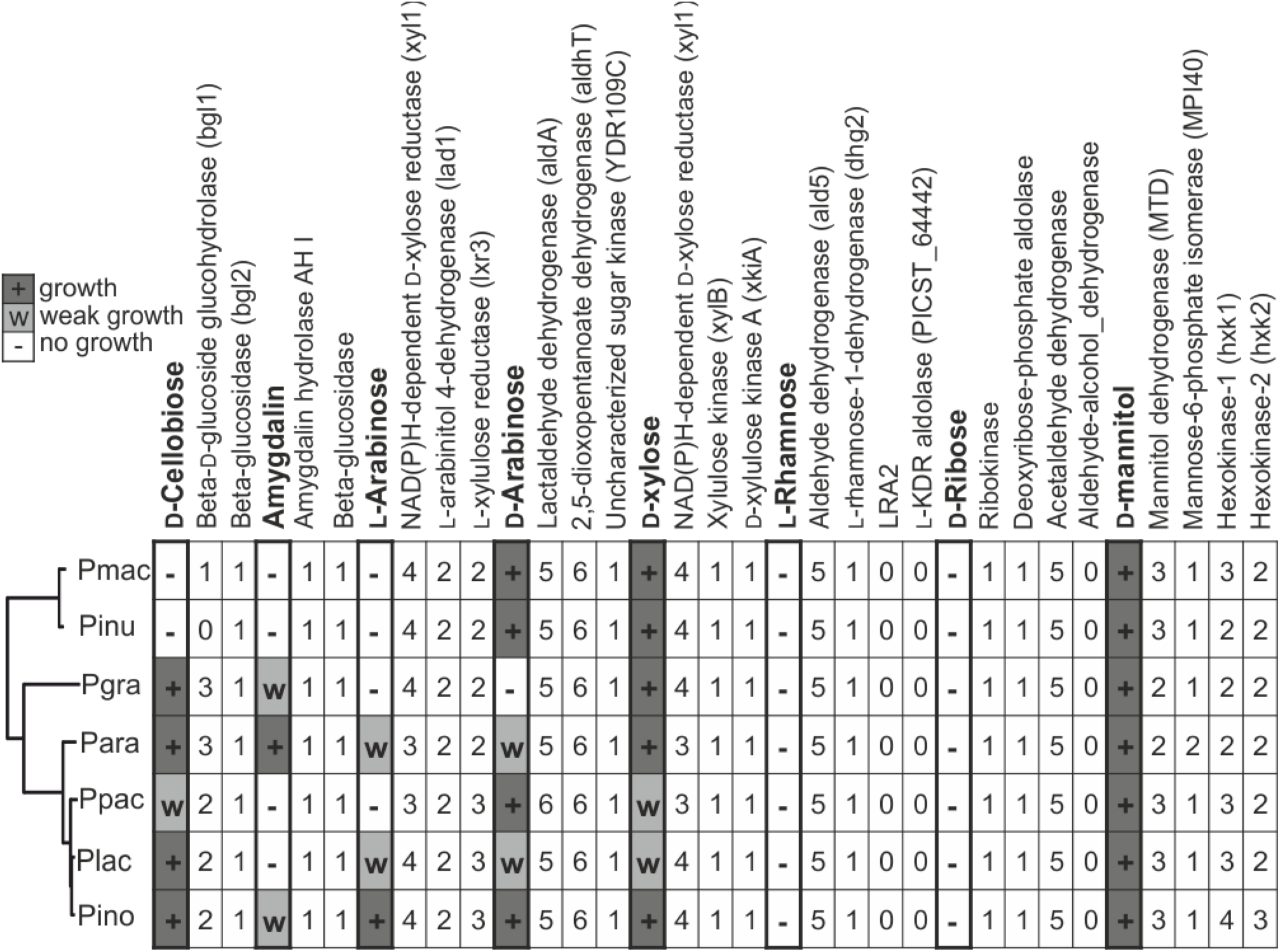
Carbon assimilation traits versus nuclear genes. Distribution of carbon assimilation patterns and related enzymes in the genus *Protomyces*. Yeast cells were grown in api^®^ 50 CH strips with a starting OD = 0.1, and were visually scored for growth on day seven (Table 4). Growth patterns were classified as growth (+), weak growth (w) and no growth (-). Numbers indicate the enzyme hits found for each carbon assimilation pathway. Genes encoding *Protomyces* carbon metabolism enzymes were identified using sequences of conserved homologs that have been characterized in model yeast species, see supplemental file 2 for the protein sequences used as BLAST queries. Selection criteria for BLAST results is an E value < 1e^−30^. Species name abbreviations used: *Protomyces inundatus* (Pinu), *P. macrosporus* (Pmac), *P. gravidus* (Pgra), *P. lactucaedebilis* (Plac), *P. inouyei* (Pino), *P. pachydermus* (Ppac), and *P. arabidopsidicola* strain C29 (Para).

**Table 4.** Carbon assimilation patterns of *Protomyces* species. Carbon assimilation was tested using 50 CH strips (biomerieuxdirect.com). Key to growth symbols: +, positive, w, weakly positive, -, negative. Pgra: *P. gravidus*, Pino: *P. inouyei*, Pinu: *P. inundatus*, Plac: *P. lactucaedebilis*, Pmac: *P. macrosporus*, Ppac: *P. pachydermus*, SC29: *P*. sp. SC29. Carbon sources listed in bold text are used differentially by these *Protomyces* species.

| Carbon source | Pgra | Pino | Pinu | Plac | Pmac | Ppac | SC29 |
| --- | --- | --- | --- | --- | --- | --- | --- |
| Glycerol | + | + | + | + | + | + | + |
| Erythritol | - | - | - | - | - | - | - |
| **D-Arabinose** | - | + | + | **w** | + | + | **W** |
| **L-Arabinose** | - | + | - | **w** | - | - | **W** |
| D-Ribose | - | - | - | - | - | - | - |
| **D-Xylose** | + | + | + | **w** | + | **w** | + |
| L- Xylose | - | - | - | - | - | - | - |
| **D-Adonitol** | - | + | - | - | - | - | - |
| Methyl-βD- xylopyranoside | - | - | - | - | - | - | - |
| D-Galactose | - | - | - | - | - | - | - |
| D-Glucose | + | + | + | + | + | + | + |
| D-Fructose | + | + | + | + | + | + | + |
| D-Mannose | + | + | + | + | + | + | + |
| L-Sorbose | - | - | - | - | - | - | - |
| **L-Rhamnose** | - | + | - | - | - | - | - |
| Dulcitol | - | - | - | - | - | - | - |
| **Inositol** | - | + | - | - | - | - | - |
| D-Mannitol | + | + | + | + | + | + | + |
| **D-Sorbitol** | - | **w** | - | + | **w** | **w** | + |
| **Methyl-αD-Mannopyranoside** | - | + | - | - | - | - | **w** |
| **Methyl-αD-Glucopranoside** | - | **w** | + | + | - | **w** | **w** |
| **N-Acetylglucosamine** | - | + | - | - | - | - | - |
| **Amygdalin** | **w** | **w** | - | - | - | - | + |
| **Arbutin** | **w** | + | - | - | - | - | + |
| **Esculin ferric citrate** | + | + | - | + | - | - | + |
| **Salicin** | **w** | + | - | - | - | - | + |
| **D-Cellobiose** | + | + | - | + | - | **w** | + |
| D-Maltose | + | + | + | + | + | + | + |
| D-Lactose | - | - | - | - | - | - | - |
| D-Melibiose | - | - | - | - | - | - | - |
| D-Saccharose (sucrose) | + | + | + | + | + | + | + |
| D-Trehalose | + | + | + | + | + | + | + |
| **Inulin** | + | + | + | + | - | **w** | - |
| **D-Melezitose** | + | + | + | + | + | + | **w** |
| D-Raffinose | + | + | + | + | + | + | + |
| Amidon (starch) | + | + | + | + | + | + | + |
| **Glycogen** | **w** | - | + | - | - | **w** | - |
| **Xylitol** | **w** | + | + | + | + | + | + |
| **Gentiobiose** | + | + | – | + | – | – | + |
| D-Turanose | + | + | + | + | + | + | + |
| **D-Lyxose** | **w** | **w** | + | – | + | **w** | – |
| D-Tagatose | – | – | – | – | – | – | – |
| D-Fucose | – | – | – | – | – | – | – |
| L-Fucose | – | – | – | – | – | – | – |
| **D-Arabitol** | + | + | – | + | + | + | + |
| **L-Arabitol** | **w** | **w** | + | – | – | – | – |
| Potassium gluconate | – | – | – | – | – | – | – |
| **Potassium 2-Ketogluconate** | **w** | + | – | + | **w** | – | **w** |
| **Potassium 5-Ketogluconate** | – | – | – | – | – | **w** | – |

**Table 5.** Diagnostic examination of *Protomyces* species. Diagnosis is based on carbon assimilation patterns from D-adonitol, Methyl-αD-mannopyranoside, L-arabinose, salicin, D-cellobiose and inulin. Carbon assimilation was performed with api^®^ CH 50 strips according to the manufacturer’s instructions. Species name abbreviations used: *Protomyces inundatus* (Pinu), *P. macrosporus* (Pmac), *P. gravidus* (Pgra), *P. lactucaedebilis* (Plac), *P. inouyei* (Pino), *P. pachydermus* (Ppac), and *P. arabidopsidicola* strain C29 (Para).

|  |  |
| --- | --- |
| 1. a) D-adonitol is assimilated | Pino |
| b) D-adonitol is not assimilated | 2 |
| 2. a) Methyl-αD-mannopyranoside is assimilated | Para |
| b) Methyl-αD-mannopyranoside is not assimilated | 3 |
| 3. a) L-arabinose is assimilated | Plac |
| b) L-arabinose is not assimilated | 4 |
| 4. a) Salicin is assimilated | Pgra |
| b) Salicin is not assimilated | 5 |
| 5. a) D-cellobiose is assimilated | Ppac |
| b) D-cellobiose is not assimilated | 6 |
| 6. a) Inulin is assimilated | Pinu |
| b) Inulin is not assimilated | Pmac |

### Phylogenomics of the genus Protomyces

The genus *Protomyces* resides within the Ascomycete subphylum Taphrinomycotina, Class Taphrinomycetes, order Taphrinales, and family Protomycetaceae. The placement of the genus *Protomyces* has been problematic since its discovery. Relationships between genera are not well supported within the subphylum Taphrinomycotina (Kurtzman et al. 2011). A well supported genome-wide phylogenetic tree constructed using both the maximum-likelihood and Bayesian inference methods with concatenated single-copy conserved proteins from species representing each family within the Taphrinomycotina, confirmed the placement of *Protomyces* within this subphylum (Fig. 6), as well as suggesting novel relationships between other genera within the subphylum.

**Fig. 6.**
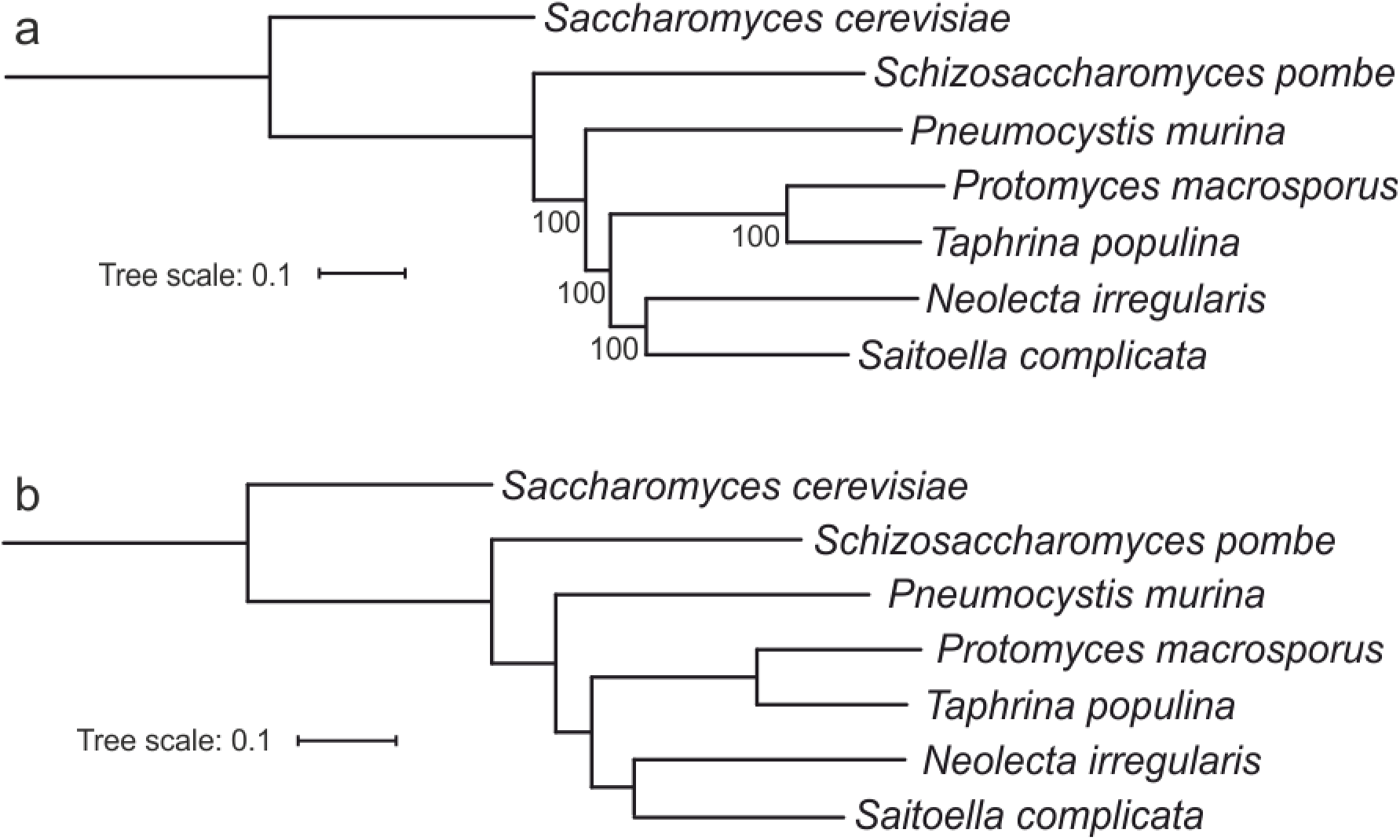
Phylogeny of the subphylum Taphrinomycotina. a), Maximum likelihood and b), Bayesian phylogenetic trees of representative species in the subphylum Taphrinomycotina. Trees were built using 636 single-copy protein sequences that were common to all species used. Alignment quality control of single-copy conserved proteins was achieved by applying sequence scores >= 0.8 in MAFFT analysis using Guidance2. *Saccharomyces cerevisiae* was used as an outgroup. Multiple aligned sequences of each species were concatenated into a single long sequence using FASconCAT_V1.0. a), the randomized axelerated maximum likelihood method (RAxML) and rapid Bootstrapping (100x) were applied for constructing the maximum-likelihood phylogenetic tree. Bootstrap support values (%) are shown at each node. b), Bayesian inference method results utilized the MrBayes software package. General time reversible (GTR) model and invgamma were chosen. Two independent analyses were started simultaneously, and 10000 generations and four chains were set for analysis run. The output files were viewed with online tool iTOL.

### P. inouyei and P. lactucaedebilis

Previous treatments of the *Protomyces* (Kurtzman et al. 2011), which utilized the same six species we have previously sequenced and analysed here, were unable to definitively conclude that these species were all genetically distinct. Our results (Table 3) clearly demonstrate that most are distinct species. However, the two species, *P. inouyei* and *P. lactucaedebilis*, were not distinct given their >96 % ANI and >97 % AAI values at the whole genome level (Table 3). The evolutionary distance between Pino and Plac is small in the phylogenetic trees built with all makers tested (Fig. 3-4). Additionally, comparisons of genomic assemblies indicated a very high level of synteny between the genomes of *P. inouyei* and *P. lactucaedebilis* (Fig. 7). Based on these data, we propose that *P. inouyei* and *P. lactucaedebilis* should be considered strains of the same species, rather than distinct species.

**Fig. 7.**
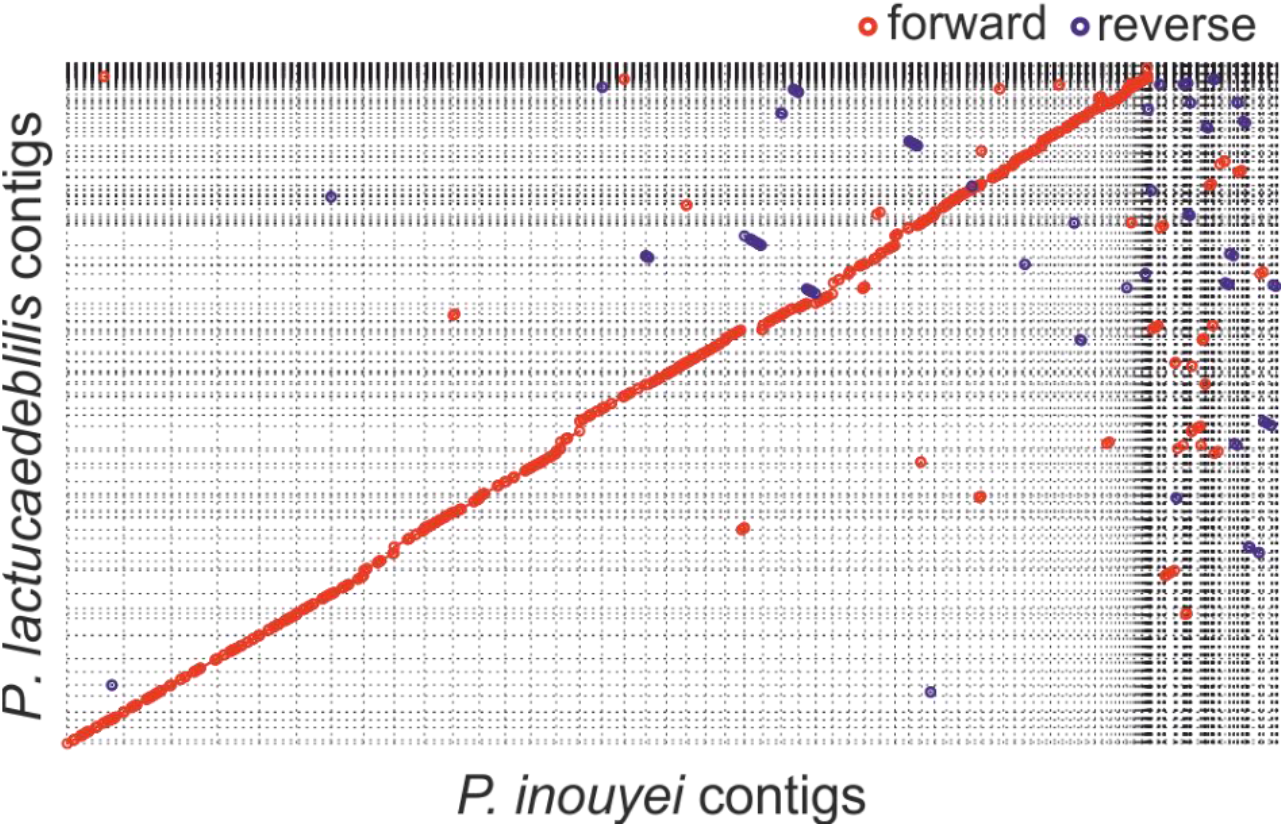
Genome synteny of *Protomyces inouyei* and *P. lactucaedebilis*. Dot-plots of whole genome alignment between *Protomyces inouyei* and *P. lactucaedebilis* contigs. Mummer 4.0.0 beta2 (nucmer) was applied for genome assembly comparison with default settings. Forward alignment is shown in red and reverse alignment in blue. Optimal co-linear order of contigs was shaped with mummerplot (parameter --fat). Mummerplot output ps files were viewed and edited with CorelDraw 2018.

## DISCUSSION

### The identification of *Protomyces arabidopsidicola sp. nov*

Based on the phylogenetic data (Fig. 3-4, and (Wang et al. 2019)) and physiological characters (Fig. 2, Tables 2 and 4), its association with *Arabidopsis* (Agler et al. 2016; Wang et al. 2019), as well as low ANI and AAI values between SC29 and other known *Protomyces* species (Table 3), we propose SC29 represents a novel species, *Protomyces arabidopsidicola sp. nov*. This yeast was isolated in an effort to establish experimental systems to study the genetics of plant yeast interactions using the genetic model plant *Arabidopsis*, for which associated yeasts were previously few to unknown (Wang et al. 2016; Wang et al. 2019). We propose the naming of this species upon the examination of a single strain. We argue that this is justified based on the strength of the phylogenetic data supporting it as a novel species, the novelty of this species that interacts with a plant widely outside the typical host range of other *Protomyces* spp., the strength of our previous data supporting its association with *Arabidopsis* (Agler et al., 2016; Wang et al. 2016; Wang et al. 2019), and the need to stimulate further research on this grossly understudied genus.

Previous phylogenetic analysis of *Protomyces* spp., used the host tissue, in which ascogenous cells were formed, as a characteristic for classification. Further, morphology and cell size comparisons were typically done with both the ascogenous cells, which only exist during natural host infection, and cultured yeast phase cells. For reasons discussed below, SC29 was not expected to cause disease, which we confirmed in controlled infection experiments (Wang et al. 2019). In the absence of infected host tissue it was not possible to obtain ascogenous cells; thus only yeast cell sizes were compared to the six *Protomyces* reference spp. Our measurements of yeast cell sizes in the reference species (Table 2) were in agreement with previously published size ranges (Kurtzman 2011).

The application of molecular tools has extensively widened our knowledge of fungal diversity and phylogeny (Blackwell 2011; Crous et al. 2015; Rosling et al. 2011). Our results with ITS and D1/D2 did not resolve all the currently studied *Protomyces* species in well supported phylogenetic trees (Fig. 3 a, b). These results suggest that the true diversity of the genus *Protomyces* is not captured or supported in the D1/D2 or ITS phylogenies. Similar results were obtained in analyses using a different set of species in our previous work (Wang et al. 2019). Also, the same conclusion have been made previously in the sister genus *Taphrina* (Rodrigues and Fonseca 2003). Our full genome based phylogenetic tree resolved all species and was well supported (Fig. 3). However, full genome sequencing is not practical for the rapid identification of *Protomyces* spp., prompting us to test a collection of five single gene nuclear markers, either individually or concatenated (Fig. 4), as in (Stielow et al. 2015). All markers performed reasonably well at species resolution and one marker, *act1*, exhibited the same architecture as the phylogenetic tree constructed with genome-wide concatenated protein data (Fig. 3, e and f; Fig. 4). Further studies will be required to test how robust *act1* is in mimicking the topology of the genome wide tree when other species are added. However, as *act1* is a commonly used marker, we propose that once a new strain has been placed in the *Protomyces* by ITS or D1/D2 sequencing, the *act1* gene sequence can be used as a secondary marker for species identification.

Carbon source utilization remains a useful tool for rapid species identification (Kurtzman et al. 2011), especially in a genus such as *Protomyces*, with a wealth of older literature and little molecular data available for comparison. Generally, the molecular underpinnings of carbon assimilation remain understudied. The availability of whole genome sequencing data offer an opportunity to correlate genomic carbon metabolism gene content with metabolic traits. Our data indicate that some traits correlate well with genomic content, while others, such as amygdalin, L-arabinose, D-arabinose, and D-xylose, genes were present for their utilization although some species were growth negative on these carbon sources (Fig. 5). A similar finding has been previously reported for D-xylose in a wider sampling of yeast species (Riley et al. 2016). The reasons for these differences remain unknown but this may account for discrepancies in carbon use traits observed between different labs or variable results seen within a single lab. Differences in the expression or conditional expression of carbon utilization genes may account for this; such latent metabolic capability has been previously suggested in a study using a wide selection of different yeasts of biotechnological interest (Riley et al. 2016).

### The Association of *Protomyces* with *Arabidopsis thaliana*

*Arabidopsis* belongs to the *Brassicaceae* (*Cruciferae*) plant family that is very distantly related to the typical *Protomyces* hosts. Thus, we sought multiple lines of evidence to support the validity of the *Protomyces-Arabidopsis* interaction. SC29 was one of three *Protomyces* strains isolated from the leaf surface of two distinct *Arabidopsis* plants at two distinct locations within the collection sites in Helsinki, Finland (Wang et al. 2016). *Protomyces* OTUs were also found over multiple years in two separate cities in Germany (Wang et al. 2019; Agler et al. 2016). All previously examined *Protomyces* spp. are heterothallic and thus require conjugation with a partner of the opposite mating type prior to transitioning into the pathogenic hyphal form. Analysis of the MAT loci from our *Protomyces* genome sequencing data (Wang et al. 2019) confirmed heterothallism; all *Protomyces* species except one had a single MAT locus with either matP or matM. Exceptionally, *P. inundatus* had two MAT loci, one bearing matP and the other matM, suggesting possible primary homothallism in this species (Wang et al. 2019). As SC29 has a single mat locus bearing only matP it is not expected to be pathogenic without another strain bearing matM. We confirmed the lack of pathogenicity in *Arabidopsis* infection experiments under a wide variety of conditions, including growth chamber experiments and overwinter field experiments, which revealed no disease symptoms (Wang et al. 2019). SC29 was then used to explore the role of phyllosphere residency in the *Protomyces* lifecycle. SC29 could persist in the *Arabidopsis* phylloplane of both sterile *in vitro* grown and soil grown plants, while titres of its closest relative *P. inouyei* rapidly decreased (Wang et al. 2019). Furthermore, SC29 could be re-isolated from *Arabidopsis* after overwintering for six months in field infection experiments (Wang et al. 2019). ITS metagenomics experiments have revealed *Protomyces* strains on the leaf surface of other plants that are not members of the Compositae or Umbelliferae plant families (Wang et al. 2016; Prior et al. 2017; Wang et al. 2019). This suggests that *Protomyces* may exploit *Arabidopsis* and possibly also multiple other alternate hosts as a phyllosphere resident. We cannot at this time exclude the possibility that SC29 is pathogenic on a different currently unknown host species or the possibility that it is pathogenic on *Arabidopsis*. The collection of more strains of *Protomyces* from *Arabidopsis* and additional experimental evidence will be necessary for a deeper understanding of SC29/ *P. arabidopsidicola* ecology.

Finally, comparative genomic analysis revealed the genomic signatures in SC29 consistent with a host jump or lifestyle change leading us to hypothesize that SC29 may have recently jumped hosts (Wang et al. 2019). Taken together, these data support that SC29 is associated with *Arabidopsis*, whose phylloplane it can utilize as a growth space, as a possible host, or alternate host.

### The genus Protomyces is not strictly related to hosts in the Compositae or Umbelliferae

The genus *Protomyces* has been narrowly defined based on the following key criteria; morphological characteristics, host plant phylogeny, and their localization with the tissues of the host (Reddy and Kramer 1975; Kurtzman 2011). Currently all *Protomyces* spp. are known to be plant pathogens, infecting hosts belonging only in the Umbelliferae (Apiaceae) and Compositae (Asteraceae) plant families. Previously, many yeasts have been excluded from the *Protomyces* based on their atypical cell sizes and association with hosts in other plant families (Reddy and Kramer 1975). Our results place SC29 as a district species within the *Protomyces* and is associated with the *Arabidopsis* phylloplane, prompting us to propose the name *P. arabidopsidicola*. These results indicate that the *Protomyces* may include species with a non-pathogenic phyllosphere resident lifestyle on alternate hosts and/or those associated with hosts outside of the Umbelliferae and Compositae, i.e. species that do not adhere to the narrow criteria previously used to define the genus. The isolation and characterization of *P. arabidopsidicola* as well as the identification of OTUs that belong to the *Protomyces* based on their ITS sequence and reside on other plant families (Wang et al. 2019) both support this view. However, as stated above, it is possible that SC29 is pathogenic on an as yet unknown host. Nonetheless, the occurrence of a *Protomyces* species in the phylloplane of a host species outside of the usual host range is a novel finding. This suggests that *Protomyces* spp. may also survive via the utilization of the phylloplane of alternate hosts. Further studies will be required to fully resolve this issue, however, we propose here that the definition of the genus *Protomyces* be broadened to allow species with a phylloplane resident lifestyle and also species associated with hosts outside of the Umbelliferae (Apiaceae) and Compositae (Asteraceae) plant families.

### Phylogenetic implications of Protomyces spp. genome sequencing

Phylogenomics with 636 conserved single copy concatenated nuclear encoded proteins confirmed the placement of *Protomyces* within the Taphrinomycotina (Fig. 6). However, our data place both *Saitoella* and *Neolecta* in a sister clade to the *Taphrina* and *Protomyces* suggesting that *Saitoella* is outside the family Protomycetaceae and order Taphrinales, where it was previously assigned (Sugiyama et al. 2006). This suggests that *Saitoella* should either define its own family to be created and named or that this genus should be assigned to the Neolectales. *Pneumocystis*, representative of the family Pneumocystidales, was previous a sister group with *Schizosaccharomyces*, but our results now suggest it may reside as outgroup between the Taphrinales and the Schizosaccharomycetales families. Further studies with a wider selection of representative species will be required to better resolve the relationships within the subphylum Taphrinomycotina.

Our results and those of many others (Liu et al. 2009; Rosling et al. 2011) suggest there are a large number of undiscovered and lost species in the Taphrinomycotina, whose discovery and analysis would aid in resolving the relationships in this fascinating subphylum. The family Protomycetaceae contains the genera *Burenia, Protomyces, Protomycopsis, Saitoella, Taphridium*, and *Volkartia*. The borders between these genera also remain poorly defined and all but *Saitoella* have similar plant pathogenic lifestyles. Unfortunately, with the exception of the genera *Protomyces* and *Saitoella*, strains and DNA sequences are not available for species in any of these genera.

Based on genomic data (Fig. 3, 4, 7 and Table 3), we propose that *P. inouyei* and *P. lactucaedebilis* are the same species. Thus, with the older name *P. inouyei*, described by Hennings in 1902, taking precedence over *P. lactucaedebilis*, described by Sawada in 1925, we propose a combining these strains under the name *P. inouyei*. These two should be differentiated by distinct strain names pending resolution of their host specificities.

Although *P. inouyei* and *P. lactucaedebilis* are thought to have distinct host specificities, infecting *Crepis* spp. and *Lactuca debilis*, respectively, this has never been formally tested by reciprocal infections. Such experiments will be required to differentiate between the two possibilities below. The new combined *P. inouyei* may be a case of different strains of the same species with distinct host specificities (*formae speciales* or subspecies). Alternatively, *P. inouyei* may be a species with a broad host range, infecting multiple host plants. The latter is perhaps the more likely possibility, as this has been seen for several other *Protomyces* species, such as *P. gravidus, P. macrosporus, P. pachydermus*, and *P. inundatus* (Table 1).

## CONCLUSIONS

The *Protomyces* sp. strain C29 isolated from the phylloplane of *Arabidopsis* is a novel species here named as *Protomyces arabidopsidicola* sp. nov. Given the novel lifestyle and the association of this new species with a plant species outside of the previously accepted host range of the members of the genus *Protomyces*, we propose that the definition of the genus is widened to include non-pathogenic phylloplane-resident species and species associated with hosts outside of the plant families Umbelliferae (Apiaceae) and Compositae (Asteraceae). Finally, we propose that the species *P. inouyei* and *P. lactucaedebilis* are not distinct species and should be merged under the name *P. inouyei*.

## TAXONOMY

### *Protomyces arabidopsidicola* Wang & Overmyer, *sp. nov*

MycoBank MB 830646.

#### Etymology

The epithet refers to the host plant (*Arabidopsis thaliana*), from which the fungus was isolated (“the *Arabidopsis-inhabiting Protomyces*”.)

#### Type

Finland, Helsinki (Lat: 60.23270N Lon: 25.06191E), leaf wash of healthy wild growing *Arabidopsis thaliana*, May 2013, K. Wang and Overmyer. (University of Helsinki HAMBI Microbial Culture Collection HAMBI3697 – holotype; The German Collection of Microorganisms and Cell Cultures GmbH, DSM 110145 – ex-type culture)

#### Description

Haploid cells of *Protomyces arabidopsidicola* sp. nov. are yeast-like and oval in shape, ranging from 2.0–12.4 μm in length and 1.4–5.6 μm in width when cultured in GYP plant for three days. Single colonies are circular and convex, yellowish in colour and become slight pinkish after about a week. Growth does not occur when temperature exceeds 30 °C or is below 8 °C when assayed on GYP agar medium for seven days; however, slow growth with colony appearance at ≽ two weeks has been observed at 4 °C. Yeast-like cells can assimilate glycerol, D-xylose, D-glucose, D-fructose, D-mannose, D-mannitol, D-sorbitol, amygdalin, arbutin, esculin ferric citrate, salicin, D-cellobiose, D-maltose, D-saccharose (sucrose), D-trehalose, D-raffinose, amidon (starch), xylitol, gentiobiose, D-turanose, D-arabitol, and weakly growth with D-arabinose, L-arabinose, Methyl-α-D-mannopyranoside, Methyl-α-D-glucopranoside, D-melezitose, Potassium 2-ketogluconate, but do not assimilate erythritol, D-ribose, L-xylose, D-adonitol, Methyl-β-D-xylopyranoside, D-galactose, L-sorbose, L-rhamnose, dulcitol, inositol, N-acetylglucosamine, D-lactose, D-melibiose, inulin, glycogen, D-lyxose, D-tagatose, D-fucose, L-fucose, L-arabitol, potassium gluconate or potassium 5-ketogluconate.

The genome size of SC29 is 11.9 Mbp (50.9 % GC content), with 5514 annotated protein-coding genes (Wang et al. 2019).

## ABBREVIATIONS

Para: *Protomyces arabidopsidicola* sp. nov.
SC29: Para strain C29
Pgra: *P. gravidus*
Pino: *P. inouyei*
Pinu: *P. inundatus*
Plac: *P. lactucaedebilis*
Pmac: *P. macrosporus*
Ppac: *P. pachydermus*

## SUPPLEMENTAL FILES

**Supplemental file 1 Queried yeast culture collections.** The file contains a list of the thirty major yeast culture collections that were queried for availability of strains for species belonging to the genus *Protomyces*.

**Supplemental file 2 Carbon utilization enzyme protein sequences.** The file contains the characterized protein sequences from model yeast species that were used as BLAST queries against the genomes of *Protomyces* spp. to identify genes involved in the utilization of various carbon sources.

## ACCESSION NUMBERS

The datasets supporting the conclusions of this article are available in the repositories listed below: Gene and genome sequences were deposited in GenBank, www.ncbi.nlm.nih.gov/ genbank, under the following accession numbers: For ITS sequences: *Protomyces arabidopsidicola* sp. nov strain SC29 (Para), LT602858, *P. gravidus* (Pgra), MK937055; *P. inouyei* (Pino), MK937056; *P. inundatus* (Pinu), MK937057; *P. lactucaedebilis* (Plac), MK937058; *P. macrosporus* (Pmac), MK937059; *P. pachydermus* (Ppac), MK937060. For D1/D2 sequences: Para, MK934482. For act1: Para, MN031257; Pgra, MN031251; Pino, MN031252; Pinu, MN031253; Plac, MN031254; Pmac, MN031255; Ppac MN031256. For rbp1: Para, MN270968; Pgra, MN270962; Pino, MN270963; Pinu, MN270964; Plac, MN270965; Pmac, MN270966; Ppac MN270967. For rbp2: Para, MN313889; Pgra, MN313883; Pino, MN313884; Pinu, MN313885; Plac, MN313886; Pmac, MN313887; Ppac MN313888. For tef1: Para, MN304745; Pgra, MN304739; Pino, MN304740; Pinu, MN304741; Plac, MN304742; Pmac, MN304743; Ppac MN304744. For tub2: Para, MN178303; Pgra, MN178297; Pino, MN178298; Pinu, MN178299; Plac, MN178300; Pmac, MN178301; Ppac, MN178302. The GenBank accession number of *Protomyces arabidopsidicola* strain C29 genome is QXMI00000000 and genome raw data in SRA is SRR8109439. Genome annotations available at genomevolution.org/coge/GenomeInfo.pl? with the following genome IDs: Para, 53653; Pgra, 53651; Pino, 53654; Pinu, 53676; Plac, 54947; Pmac, 53670; Ppac 54948. The strain SC29 has been deposited as the holotype in the University of Helsinki Microbial Domain Biological Resource Centre (HAMBI) Culture Collection, www.helsinki.fi/en/infrastructures/biodiversity-collections/infrastructures/microbial-domain-biological-resource-centre-hambi, under the accession no. HAMBI3697, and an isotype culture was deposited in the Collection of Microorganisms and Cell Cultures (DSMZ) culture collection, www.dsmz.de, under the accession no. DSM 110145. The species name *Protomyces arabidopsidicola* has been registered with Mycobank, http://www.mycobank.org/, under the accession no. MB 830646.

## AUTHOR CONTRIBUTIONS

KO, TS, and KW designed experiments, TS and KW performed experiments, KW performed all bioinformatics analyses. KW and KO, wrote the manuscript, all authors edited and approved the manuscript.

## ACKNOWLEDGEMENTS

Computing resources provided by the Finnish IT Center for Science (CSC; www.csc.fi) are gratefully acknowledged. We wish to thank Prof. Dr. Dominik Begerow and Prof. Daniel Croll for their critical comments on this manuscript during the external examination of Kai Wang’s PhD thesis. We also would like to thank Prof. Dr. Michael Weiss and Dr. Konstanze Bensch for their guidance concerning the naming of *P. arabidopsidicola*.

## REFERNCES

Agler MT, Ruhe J, Kroll S, Morhenn C, Kim S-T, Weigel D, Kemen EM (2016) Microbial hub taxa link host and abiotic factors to plant microbiome variation. PLoS biology 14 (1):e1002352. doi:10.1371/journal.pbio.1002352.

Bacigálová K (2008) *Protomyces buerenianus* (Protomycetaceae)—a new species for Slovakia. Biologia 63 (1): 40–43.

Bacigálová K, Piątek M, Wołkowycki M (2005) *Protomyces cirsii-oleracei* (Fungi, Protomycetales), a new species for Poland. Polish Botanical Journal 50 (1):77–82.

Bary A, Garnsey HEF (1887) Comparative morphology and biology of the fungi, mycetozoa and bacteria, vol 2. Clarendon press, Oxford.

Blackwell M (2011) The Fungi: 1, 2, 3… 5.1 million species? American Journal of Botany 98 (3):426–438.

Boundy - Mills KL, Glantschnig E, Roberts IN, Yurkov A, Casaregola S, Daniel HM, Groenewald M, Turchetti B (2016) Yeast culture collections in the twenty - first century: New opportunities and challenges. Yeast 33 (7):243–260.

Crous PW, Hawksworth DL, Wingfield MJ (2015) Identifying and naming plant-pathogenic fungi: past, present, and future. Annual Review of Phytopathology 53:247–267.

Emms DM, Kelly S (2015) OrthoFinder: solving fundamental biases in whole genome comparisons dramatically improves orthogroup inference accuracy. Genome biology 16 (1):157.

Hibbett DS, Binder M, Bischoff JF, Blackwell M, Cannon PF, Eriksson OE, Huhndorf S, James T, Kirk PM, Lücking R (2007) A higher-level phylogenetic classification of the Fungi. Mycological Research 111 (5):509–547.

James TY, Kauff F, Schoch CL, Matheny PB, Hofstetter V, Cox CJ, Celio G, Gueidan C, Fraker E, Miadlikowska J (2006) Reconstructing the early evolution of Fungi using a six-gene phylogeny. Nature 443 (7113): 818.

Kurtzman C (1993) The systematics of ascomycetous yeasts defined from ribosomal RNA sequence divergence: theoretical and practical considerations. In: Reynolds DR, Taylor JW (eds) The Fungal Holomorph: Mitotic, Meiotic Pleomorphic Speciation in Fungal Systematics. CAB International, Wallingford, pp 271–279.

Kurtzman CP (2011) *Protomyces* Unger (1833). In: Kurtzman CP, Fell JW, Boekhout T (eds) The yeasts: a taxonomic study. 5th edn. Elsevier, Amsterdam, pp 725–731.

Kurtzman CP, Fell JW, Boekhout T (2011) The yeasts: a taxonomic study. 5th edn. Elsevier, Amsterdam.

Kurtzman CP, Robnett CJ (1998) Identification and phylogeny of ascomycetous yeasts from analysis of nuclear large subunit (26S) ribosomal DNA partial sequences. Antonie van Leeuwenhoek 73 (4):331–371.

Kurtzman CP, Sugiyama J (2015) Saccharomycotina and Taphrinomycotina: The Yeasts and Yeastlike Fungi of the Ascomycota. In: McLaughlin DJ, Spatafora JW (eds) The mycota VII: Systematics and evolution: Part B. 2nd Edition edn. Springer Berlin Heidelberg, Berlin, Heidelberg, pp 3–33. doi: 10.1007/978-3-662-46011-5_1.

Larkin MA, Blackshields G, Brown N, Chenna R, McGettigan PA, McWilliam H, Valentin F, Wallace IM, Wilm A, Lopez R (2007) Clustal W and Clustal X version 2.0. Bioinformatics 23 (21):2947–2948.

Letunic I, Bork P (2016) Interactive tree of life (iTOL) v3: an online tool for the display and annotation of phylogenetic and other trees. Nucleic Acids Research 44 (W1):W242–W245.

Liu Y, Leigh JW, Brinkmann H, Cushion MT, Rodriguez-Ezpeleta N, Philippe H, Lang BF (2009) Phylogenomic analyses support the monophyly of taphrinomycotina, including *Schizosaccharomyces* fission yeasts. Molecular Biology and Evolution 26 (1):27–34. doi:10.1093/molbev/msn221.

Lohwag H (1934) Zu Lycoperdellon. Annales Mycologici 32: 244–255.

Mix A (1924) Biological and cultural studies of *Exoascus deformans*. Phytopathology 14:217–233.

Nishida H, Hamamoto M, Sugiyama J (2011) Draft genome sequencing of the enigmatic yeast *Saitoella complicata*. The Journal of General and Applied Microbiology 57 (4):243–246.

Nishida H, Sugiyama J (1994) Archiascomycetes: detection of a major new lineage within the Ascomycota. Mycoscience 35 (4):361–366.

Nishida H, Sugiyama J (1995) A common group I intron between a plant parasitic fungus and its host. Molecular Biology and Evolution 12 (5):883–886.

Piepenbring M, Bauer R (1997) Erratomyces, a new genus of Tilletiales with species on Leguminosae. Mycologia 89 (6):924–936.

Prior R, Mittelbach M, Begerow D (2017) Impact of three different fungicides on fungal epi-and endophytic communities of common bean *(Phaseolus vulgaris)* and broad bean *(Vicia faba)*. Journal of Environmental Science and Health, Part B 52 (6):376–386.

Reddy MS, Kramer C (1975) A taxonomic revision of the Protomycetales. Mycotaxon 3 (1):1–50.

Riley R, Haridas S, Wolfe KH, Lopes MR, Hittinger CT, Göker M, Salamov AA, Wisecaver JH, Long TM, Calvey CH (2016) Comparative genomics of biotechnologically important yeasts. Proceedings of the National Academy of Sciences, USA 113 (35):9882–9887.

Rodrigues MG, Fonseca Á (2003) Molecular systematics of the dimorphic ascomycete genus *Taphrina*. International journal of systematic and evolutionary microbiology 53 (2):607–616.

Rodriguez-R LM, Konstantinidis KT (2016) The enveomics collection: a toolbox for specialized analyses of microbial genomes and metagenomes. PeerJ Preprints. doi: 10.7287/peerj.preprints.1900v1.

Ronquist F, Teslenko M, Van Der Mark P, Ayres DL, Darling A, Höhna S, Larget B, Liu L, Suchard MA, Huelsenbeck JP (2012) MrBayes 3.2: efficient Bayesian phylogenetic inference and model choice across a large model space. Systematic Biology 61 (3):539–542.

Rosling A, Cox F, Cruz-Martinez K, Ihrmark K, Grelet G-A, Lindahl BD, Menkis A, James TY (2011) Archaeorhizomycetes: unearthing an ancient class of ubiquitous soil fungi. Science 333 (6044):876–879.

Sadebeck R (1884) Untersuchungen iiber die Pilzgattung Exoascus. Jahrbuch der Hamburgischen Wissenschaftlichen Anstalten 1:93–124.

Stamatakis A (2014) RAxML version 8: a tool for phylogenetic analysis and post-analysis of large phylogenies. Bioinformatics 30 (9):1312–1313. doi:10.1093/bioinformatics/btu033.

Stielow JB, Levesque CA, Seifert KA, Meyer W, Iriny L, Smits D, Renfurm R, Verkley G, Groenewald M, Chaduli D (2015) One fungus, which genes? Development and assessment of universal primers for potential secondary fungal DNA barcodes. Persoonia 35: 242.

Sugiyama J, Hosaka K, Suh S-O (2006) Early diverging Ascomycota: phylogenetic divergence and related evolutionary enigmas. Mycologia 98 (6):996–1005.

Sugiyama J, Nagahama T, Nishida H (1996) Fungal diversity and phylogeny with emphasis on 18S ribosomal DNA sequence divergence. In: RR C, U S, K O (eds) Microbial diversity in time and space. Springer, Boston, pp 41–51.

Tubaki K (1957) Biological and cultural studies of three species of *Protomyces*. Mycologia 49 (1):44–54.

Turland NJ, Wiersema JH, Barrie FR, Greuter W, Hawksworth D, Herendeen PS, Knapp S, Kusber W-H, Li D-Z, Marhold K (2018) International code of nomenclature for algae, fungi, and plants (Shenzhen Code) adopted by the Nineteenth International Botanical Congress Shenzhen, China, July 2017. Regnum Vegetabile 159 Glashütten: Koeltz Botanical Books.

Unger F (1833) Die Exantheme der Pflanzen. Gerold, Wien: 421.

Walker WF (1985) 5S ribosomal RNA sequences from ascomycetes and evolutionary implications. Systematic and Applied Microbiology 6 (1):48–53.

Van Eijk G, Roeymans H (1982) Distribution of carotenoids and sterols in relation to the taxonomy of *Taphrina* and *Protomyces*. Antonie van Leeuwenhoek 48 (3):257–264.

Wang K, Sipilä T, Rajaraman S, Safronov O, Laine P, Auzane A, Mari A, Auvinen P, Paulin L, Kemen E, Salojärvi J, Overmyer K (2019) A novel phyllosphere resident *Protomyces* species that interacts with the *Arabidopsis* immune system. Biorxiv:594028. doi: 10.1101/594028%JbioRxiv.

Wang K, Sipilä TP, Overmyer K (2016) The isolation and characterization of resident yeasts from the phylloplane of *Arabidopsis thaliana*. Scientific Reports 6: 39403.

Von Arx J, Weijman A (1979) Conidiation and carbohydrate composition in some *Candida* and *Torulopsis* species. Antonie van Leeuwenhoek 45 (4):547–555.

