## Supplemental file 1 for "A novel *Arabidopsis* phyllosphere resident *Protomyces* sp. and a re-examination of genus *Protomyces* based on genome sequence data"

**Supplemental file 1. Yeast culture collections queried.** List of thirty major yeast culture collections that were queried for availability of strains of species belonging to the genus *Protomyces*. Search was done in in July 2019. The queried collections were:

The China General Microbiological Culture Collection Center, Beijing, China (CGMCC; http://www.cgmcc.net/english)

Microbial Domain Biological Resource Centre, Helsinki, Finland (HAMBI; https://www.helsinki.fi/en/infrastructures/biodiversity-collections/infrastructures/microbial-domain-biological-resource-centre-hambi).

VTT Technical Research Center of Finland Culture Collection, Espoo, Finland (http://culturecollection.vtt.fi/m).

The German Collection of Microorganisms and Cell Cultures GmbH, Braunschweig, Germany (DSMZ; www.dsmz.de).

The Industrial Yeasts Collection, Perugia, Italy (DBVPG; http://www.dbvpg.unipg.it/index.php/en).

The Japan Collection of Microorganisms, Koyadai, Tsukuba, Ibaraki, Japan (JCM; jcm.brc.riken.jp/en/).

Biological Resource Center NITE, Chiba, Japan (NBRC; www.nite.go.jp/en/nbrc).

CBS-KNAW Fungal Biodiversity Centre, Utrecht, The Netherlands (CBS; http://www.westerdijkinstitute.nl/Collections/).

The Portuguese Yeast Culture Collection, Caparica, Portugal (PYCC; http://pycc.bio-aware.com/).

The All-Russian Collection of Microorganisms, Pushchino and Moskow Russia (VKM; http://www.vkm.ru).

Culture Collection of Yeasts, Bratislava, Slovakia (CCY; http://ccy.sk/index.php/en/).

National Collections of Yeast Cultures, Norwich, UK (NCYC; www.ncyc.co.uk).

Agricultural Research Service Culture Collection, Peoria, IL, USA (NRRL; nrrl.ncaur.usda.gov).

American Type Culture Collection, Manassas, VA, USA (ATCC; www.atcc.org).

University of California Phaff Culture Collection, Davis, CA, USA (UCD-FST; http://phaffcollection.ucdavis.edu).

Belgian Coordinated Collection of Microorganisms, Louvain-la-Neuve, Belgium (BCCM/MCLU; http://bccm.belspo.be/).

Bioresource Collection and Research Center, Hsinchu, Taiwan (BCRC; catalog.bcrc.firdi.org.tw).

Chinese Center for Industrial Culture Collection, Beijing, China (CICC; http://english.china-cicc.org/).

Centre International de Ressources Microbiennes-Levures, Levure, France (CIRM-Levures; https://www6.inra.fr/cirm_eng/Yeasts).

Spanish Type Culture Collection, Valencia, Spain (CECT; https://www.uv.es/cect).

Korean Collection for Type Cultures, Jeollabuk-do, Korea (KCTC; https://kctc.kribb.re.kr/En/Kctc.aspx).

ZIM Collection of Industrial Microorganisms, University of Ljubljana, Ljubljana, Slovenia (ZIM; www.bf.uni-lj.si/zt/biotech/chair/index.html).

Universidade Federal de Pernambuco, Micoteca do Departmento de Micologia, Recife, Brazil (URM; https://www.ufpe.br/micoteca/).

Lallemand Yeast Culture Collection, Lallemand Inc., Canada (LYCC; www.lallemand.com/).

National Collection of Agricultural and Industrial Microorganisms, Budapest, Hungary (NCAIM; ncaim.etk.szie.hu/).

National Bank for Industrial Microorganisms and Cell Cultures, University of Chemical Technology and Metallurgy, Sofia, Bulgaria (NBIMCC; www.nbimcc.org).

Microbial Type Culture Collection, Institute of Microbial Technology, Chandigarh, India (MTCC; https://mtccindia.res.in/).

Collection of Industrial Microorganisms, Institute of Agricultural and Food Biotechnology, Warsaw, Poland (IAFB; https://cim.ibprs.pl/).

Food Science Australia, Ryde, CSIRO, North Ryde, Australia (FRR; www.foodscience.csiro.au/fcc/services.htm).

The UAMH Centre for Global Microfungal Biodiversity, Toronto, Canada (UAMH; www.uamh.ca).
