## Supplemental file 2 for "A novel *Arabidopsis* phyllosphere resident *Protomyces* sp. and a re-examination of genus *Protomyces* based on genome sequence data"

**Supplemental file 2. Carbon utilization enzyme protein sequences.** Characterized protein sequences from model species that were used as BLAST queries against the genomes of *Protomyces* spp. to identify genes involved in the utilization of various carbon sources.

>sp|P25553|ALDA_ECOLI Lactaldehyde dehydrogenase OS=Escherichia coli (strain K12) GN=aldA PE=1 SV=2

MSVPVQHPMYIDGQFVTWRGDAWIDVVNPATEAVISRIPDGQAEDARKAIDAAERAQPEWE
ALPAIERASWLRKISAGIRERASEISALIVEEGGKIQQLAEVEVAFTADYIDYMAEWARRYEGEI
IQSDRPGENILLFKRALGVTTGILPWNFPFFLIARKMAPALLTGNTIVIKPSEFTPNNAIAFAKIV
DEIGLPRGVFNLVLGRGETVGQELAGNPKVAMVSMTGSVSAGEKIMATAAKNITKVCLELGG
KAPAIVMDDADLELAVKAIVDSRVINSGQVCNCAERVYVQKGIYDQFVNRLGEAMQAVQFGN
PAERNDIAMGPLINAAALERVEQKVARAVEEGARVAFGGKAVEGKGYYYPPTLLLDVRQEM
SIMHEETFGPVLPVVAFDTLEDAISMANDSDYGLTSSIYTQNLNVAMKAIKGLKFGETYINREN
FEAMQGFHAGWRKSGIGGADGKHGLHEYLQTQVVYLQS

>sp|Q97U96|ARAD_SULSO Arabinonate dehydratase OS=Sulfolobus solfataricus (strain ATCC 35092 / DSM 1617 / JCM 11322 / P2) GN=araD PE=1 SV=1

MIKDIRTYKLCYEGINDERDALAIKGLAEHPMEIVATEIETSDGYVGYGESLAYGCSDAVQVTI
EKILKPLLLKEDEELIEYLWDKMYKATLRFGRRGIAIAGISGVDTALWDIMGKKAKKPIYKLLGG
SKRKVRAYITGGYYSEKKDLEKLRDEEAYYVKMGFKGIKVKIGAKSMEEDIERLKAIREVVGE
DVKIAVDANNVYTFEEALEMGRRLEKLGIWFFEEPIQTDYLDLSARLAEELEVPIAGYETAYTR
WEFYEIMRKRAVDIVQTDVMWTGGISEMMKIGNMAKVMGYPLIPHYSAGGISLIGNLHVAAA
LNSPWIEMHLRKNDLRDKIFKESIEIDNGHLVVPDRPGLGYTIRDGVFEEYKCKS

>sp|Q97UA1|KGSDH_SULSO 2,5-dioxopentanoate dehydrogenase OS=Sulfolobus solfataricus (strain ATCC 35092 / DSM 1617 / JCM 11322 / P2) GN=aldhT PE=1 SV=1

MKSYQGLADKWIKGSGEEYLDINPADKDHVLAKIRLYTKDDVKEAINKAVAKFDEWSRTPAP
KRGSILLKAGELMEQEAQEFALLMTLEEGKTLKDSMFEVTRSYNLLKFYGALAFKISGKTLPS
ADPNTRIFTVKEPLGVVALITPWNFPLSIPVWKLAPALAAGNTAVIKPATKTPLMVAKLVEVLS
KAGLPEGVVNLVVGKGSEVGDTIVSDDNIAAVSFTGSTEVGKRIYKLVGNKNRMTRIQLELG
GKNALYVDKSADLTLAAELAVRGGFGLTGQSCTATSRLIINKDVYTQFKQRLLERVKKWRVG
PGTEDVDMGPVVDEGQFKKDLEYIEYGKNVGAKLIYGGNIIPGKGYFLEPTIFEGVTSDMRLF
KEEIFGPVLSVTEAKDLDEAIRLVNAVDYGHTAGIVASDIKAINEFVSRVEAGVIKVNKPTVGLE
LQAPFGGFKNSGATTWKEMGEDALEFYLKEKTVYEGW

>sp|P0AB87|FUCA_ECOLI L-fuculose phosphate aldolase OS=Escherichia coli (strain K12) GN=fucA PE=1 SV=1

MERNKLARQIIDTCLEMTRLGLNQGTAGNVSVRYQDGMLITPTGIPYEKLTESHIVFIDGNGK
HEEGKLPSSEWRFHMAAYQSRPDANAVVHNHAVHCTAVSILNRSIPAIHYMIAAAGGNSIPC
APYATFGTRELSEHVALALKNRKATLLQHHGLIACEVNLEKALWLAHEVEVLAQLYLTTLAITD
PVPVLSDEEIAVVLEKFKTYGLRIEE

>sp|P11553|FUCK_ECOLI L-fuculokinase OS=Escherichia coli (strain K12) GN=fucK PE=1 SV=3

MLSGYIAGAIMKQEVILVLDCGATNVRAIAVNRQGKIVARASTPNASDIAMENNTWHQWSLD
AILQRFADCCRQINSELTECHIRGIAVTTFGVDGALVDKQGNLLYPIISWKCPRTAAVMDNIER
LISAQRLQAISGVGAFSFNTLYKLVWLKENHPQLLERAHAWLFISSLINHRLTGEFTTDITMAG
TSQMLDIQQRDFSPQILQATGIPRRLFPRLVEAGEQIGTLQNSAAAMLGLPVGIPVISAGHDT
QFALFGAGAEQNEPVLSSGTWEILMVRSAQVDTSLLSQYAGSTCELDSQAGLYNPGMQWL
ASGVLEWVRKLFWTAETPWQMLIEEARLIAPGADGVKMQCDLLSCQNAGWQGVTLNTTRG
HFYRAALEGLTAQLQRNLQMLEKIGHFKASELLLVGGGSRNTLWNQIKANMLDIPVKVLDDA
ETTVAGAALFGWYGVGEFNSPEEARAQIHYQYRYFYPQTEPEFIEEV

>sp|P69922|FUCI_ECOLI L-fucose isomerase OS=Escherichia coli (strain K12) GN=fucI PE=1 SV=1

MKKISLPKIGIRPVIDGRRMGVRESLEEQTMNMAKATAALLTEKLRHACGAAVECVISDTCIA
GMAEAAACEEKFSSQNVGLTITVTPCWCYGSETIDMDPTRPKAIWGFNGTERPGAVYLAAA
LAAHSQKGIPAFSIYGHDVQDADDTSIPADVEEKLLRFARAGLAVASMKGKSYLSLGGVSMGI
AGSIVDHNFFESWLGMKVQAVDMTELRRRIDQKIYDEAELEMALAWADKNFRYGEDENNKQ
YQRNAEQSRAVLRESLLMAMCIRDMMQGNSKLADIGRVEESLGYNAIAAGFQGQRHWTDQ
YPNGDTAEAILNSSFDWNGVREPFVVATENDSLNGVAMLMGHQLTGTAQVFADVRTYWSP
EAIERVTGHKLDGLAEHGIIHLINSGSAALDGSCKQRDSEGNPTMKPHWEISQQEADACLAA
TEWCPAIHEYFRGGGYSSRFLTEGGVPFTMTRVNIIKGLGPVLQIAEGWSVELPKDVHDILNK
RTNSTWPTTWFAPRLTGKGPFTDVYSVMANWGANHGVLTIGHVGADFITLASMLRIPVCMH
NVEETKVYRPSAWAAHGMDIEGQDYRACQNYGPLYKR

>sp|Q97UA0|KDAD_SULSO 2-dehydro-3-deoxy-D-arabinonate dehydratase OS=Sulfolobus solfataricus (strain ATCC 35092 / DSM 1617 / JCM 11322 / P2) GN=kdaD PE=1 SV=1

MHFIMMKLFRVVKRGYYISYAILDNSTIIRLDEDPIKALMRYSENKEVLGDRVTGIDYQSLLKSF
QINDIRITKPIDPPEVWGSGISYEMARERYSEENVAKILGKTIYEKVYDAVRPEIFFKATPNRCV
GHGEAIAVRSDSEWTLPEPELAVVLDSNGKILGYTIMDDVSARDLEAENPLYLPQSKIYAGCC
AFGPVIVTSDEIKNPYSLDITLKIVREGRVFFEGSVNTNKMRRKIEEQIQYLIRDNPIPDGTILTT
GTAIVPGRDKGLKDEDIVEITISNIGTLITPVKKRRKIT

>sp|Q04585|YDR09_YEAST Uncharacterized sugar kinase YDR109C OS=Saccharomyces cerevisiae (strain ATCC 204508 / S288c) GN=YDR109C PE=1 SV=1

MKSRKRQNNMQNETREPAVLSSQETSISRISPQDPEAKFYVGVDVGTGSARACVIDQSGNM
LSLAEKPIKREQLISNFITQSSREIWNAVCYCVRTVVEESGVDPERVRGIGFDATCSLVVVSAT
NFEEIAVGPDFTNNDQNIILWMDHRAMKETEEINSSGDKCLKYVGGQMSVEMEIPKIKWLKN
NLEAGIFQDCKFFDLPDYLTFKATGKENRSFCSAVCKQGFLPVGVEGSDIGWSKEFLNSIGL
SELTKNDFERLGGSLREKKNFLTAGECISPLDKKAACQLGLTEHCVVSSGIIDAYAGWVGTVA
AKPESAVKGLAETENYKKDFNGAIGRLAAVAGTSTCHILLSKNPIFVHGVWGPYRDVLARGF
WAAEGGQSCTGVLLDHLITTHPAFTELSHMANLAGVSKFEYLNKILETLVEKRKVRSVISLAK
HLFFYGDYHGNRSPIADPNMRACIIGQSMDNSIEDLAVMYLSACEFISQQTRQIIEVMLKSGH

EINAIFMSGGQCRNSLLMRLLADCTGLPIVIPRYVDAAVVFGSALLGAAASEDFDYTREKRTL
KGQKSSQTKTERFNDSYSSIQKLSMEDRNSTNGFVSPHNLQLSTPSAPAKINNYSLPICTQQ
PLDKTSEESSKDASLTVGQESLGEGRYNGTSFLWKVMQELTGNARIVNPNEKTHPDRILLDT
KYQIFLDMIETQRKYRRMVDKVEGSFSR

>tr|Q9I1Q0|Q9I1Q0_PSEAE Probable aldehyde dehydrogenase OS=Pseudomonas aeruginosa (strain ATCC 15692 / DSM 22644 / CIP 104116 / JCM 14847 / LMG 12228 / 1C / PRS 101 / PAO1) GN=PA2217 PE=4 SV=1

MTAILGHNFIGGARSAAGTLFLQSLDAASGEALPYRFVQATPEEVDAAAEAAASAYPHYRQL
PASRRAEFLDTIAAELDALDDDFVAIVCRETALPATRIQGERSRTSGQLRLFAEVLRRGDFHG
ARIDRARPERKPLPRVDLRQCRIGLGPVAVFGASNFPLAFSTAGGDTAAALAAGCPVVFKAH
SGHMATAERVAAAILRAAERTGMPAGVFNMIYGGGVGERLVRHPAIQAVGFTGSLKGGRAL
CDMAAARAQPIPVFAEMSSINPVVLLPAALKKRGEAVADELSASVVLGCGQFCTNPGLVIGIR
SAQFSAFLERFAARMDDQPAQTMLNTGTLASYEKGLAALHAHPRVRHLAGQPQEGRQARP
QLFQADVSLLLEGDELLQEEVFGPASVVVEVADHAELKRALHGLHGQLTATLIAEAEDLASFA
DLVPLLEQKAGRLLLNGYPTGVEVCDAMVHGGPYPATSDARGTSVGTLAIDRFLRPVCYQN
YPDAWLPEALKDGNPLGIARLVDGIVTRAAVA

>Ss-araDH_tr|Q97YM2|Q97YM2_SULSO Alcohol dehydrogenase (Zn containing) (Adh-4) OS=Sulfolobus solfataricus (strain ATCC 35092 / DSM 1617 / JCM 11322 / P2) GN=adh-4 PE=1 SV=1

MENVNMVKSKAALLKKFSEPLSIEDVNIPEPQGEEVLIRIGGAGVCRTDLRVWKGVEAKQGF
RLPIILGHENAGTIVEVGELAKVKKGDNVVVYATWGDLTCRYCREGKFNICKNQIIPGQTTNG
GFSEYMLVKSSRWLVKLNSLSPVEAAPLADAGTTSMGAIRQALPFISKFAEPVVIVNGIGGLA
VYTIQILKALMKNITIVGISRSKKHRDFALELGADYVSEMKDAESLINKLTDGLGASIAIDLVGTE
ETTYNLGKLLAQEGAIILVGMEGKRVSLEAFDTAVWNKKLLGSNYGSLNDLEDVVRLSESGKI
KPYIIKVPLDDINKAFTNLDEGRVDGRQVITP

>tr|Q88NF5|Q88NF5_PSEPK Putative alpha-ketoglutarate semialdehyde dehydrogenase OS=Pseudomonas putida (strain ATCC 47054 / DSM 6125 / NCIMB 11950 / KT2440) GN=PP_1256 PE=4 SV=1

MPLTGNLLIGQRPVTGSRDAIRAIDPTTGQTLEPAYLGGTGEHVAQACALAWAAFDAYRETS
LEQRAEFLEAIATQIEALGDALIDRAVIETGLPKARIQGERGRTCTQLRTFARTVRAGEWLDVR
IDSALPERQPLPRADLRQRQVALGPVAVFGASNFPLAFSVAGGDTASALAAGCPVVVKAHSA
HPGTSELVGQAVAQAVKQCGLPEGVFSLLYGSGREVGIALVSDPRIKAVGFTGSRSGGMAL
CQAAQARPEPIPVYAEMSSINPVFLFDAALQARAEALAQGFVASLTQGAGQFCTNPGLVIAR
QGPALQRFITAAAGYVQQGAAQTMLTPGIFSAYQAGIAALADNPHAQAITSGQAGQGPNQC
QAQLFVTQAEAFLADPALQAEVFGAASLVVACTDDEQVRQVAEHLEGQLTATLQLDEADIDS
ARALLPTLERKAGRILVNGWPTGVEVCDAMVHGGPFPATSDARTTSVGTAAILRFLRPVCYQ
DVPDALLPQALKHGNPLQLRRLLDGKRED

>sp|P08204|ARAB_ECOLI Ribulokinase OS=Escherichia coli (strain K12) GN=araB PE=1 SV=4

MAIAIGLDFGSDSVRALAVDCATGEEIATSVEWYPRWQKGQFCDAPNNQFRHHPRDYIESM
EAALKTVLAELSVEQRAAVVGIGVDSTGSTPAPIDADGNVLALRPEFAENPNAMFVLWKDHT
AVEEAEEITRLCHAPGNVDYSRYIGGIYSSEWFWAKILHVTRQDSAVAQSAASWIELCDWVP
ALLSGTTRPQDIRRGRCSAGHKSLWHESWGGLPPASFFDELDPILNRHLPSPLFTDTWTADI
PVGTLCPEWAQRLGLPESVVISGGAFDCHMGAVGAGAQPNALVKVIGTSTCDILIADKQSVG
ERAVKGICGQVDGSVVPGFIGLEAGQSAFGDIYAWFGRVLGWPLEQLAAQHPELKTQINAS
QKQLLPALTEAWAKNPSLDHLPVVLDWFNGRRTPNANQRLKGVITDLNLATDAPLLFGGLIA
ATAFGARAIMECFTDQGIAVNNVMALGGIARKNQVIMQACCDVLNRPLQIVASDQCCALGAAI
FAAVAAKVHADIPSAQQKMASAVEKTLQPCSEQAQRFEQLYRRYQQWAMSAEQHYLPTSA
PAQAAQAVATL

>sp|P08202|ARAA_ECOLI L-arabinose isomerase OS=Escherichia coli (strain K12) GN=araA PE=1 SV=3

MTIFDNYEVWFVIGSQHLYGPETLRQVTQHAEHVVNALNTEAKLPCKLVLKPLGTTPDEITAI
CRDANYDDRCAGLVVWLHTFSPAKMWINGLTMLNKPLLQFHTQFNAALPWDSIDMDFMNL
NQTAHGGREFGFIGARMRQQHAVVTGHWQDKQAHERIGSWMRQAVSKQDTRHLKVCRFG
DNMREVAVTDGDKVAAQIKFGFSVNTWAVGDLVQVVNSISDGDVNALVDEYESCYTMTPAT
QIHGKKRQNVLEAARIELGMKRFLEQGGFHAFTTTFEDLHGLKQLPGLAVQRLMQQGYGFA
GEGDWKTAALLRIMKVMSTGLQGGTSFMEDYTYHFEKGNDLVLGSHMLEVCPSIAAEEKPIL
DVQHLGIGGKDDPARLIFNTQTGPAIVASLIDLGDRYRLLVNCIDTVKTPHSLPKLPVANALWK
AQPDLPTASEAWILAGGAHHTVFSHALNLNDMRQFAEMHDIEITVIDNDTRLPAFKDALRWN
EVYYGFRR

>sp|P08203|ARAD_ECOLI L-ribulose-5-phosphate 4-epimerase AraD OS=Escherichia coli (strain K12) GN=araD PE=1 SV=2

MLEDLKRQVLEANLALPKHNLVTLTWGNVSAVDRERGVFVIKPSGVDYSVMTADDMVVVSIE
TGEVVEGTKKPSSDTPTHRLLYQAFPSIGGIVHTHSRHATIWAQAGQSIPATGTTHADYFYGT
IPCTRKMTDAEINGEYEWETGNVIVETFEKQGIDAAQMPGVLVHSHGPFAWGKNAEDAVHN
AIVLEEVAYMGIFCRQLAPQLPDMQQTLLDKHYLRKHGAKAYYGQ

>sp|Q91XV4|DCXR_MESAU L-xylulose reductase OS=Mesocricetus auratus GN=DCXR PE=1 SV=1

MDLGLAGRRALVTGAGKGIGRSTVLALQAAGAHVVAVSRTQADLDSLVSECPGVETVCVDL
ADWEATEQALSSVGPVDLLVNNAAVALLQPFLEVTKEAFDMSFNVNLRAVIQVSQIVARGMI
ARGAPGAIVNVSSQASQRALANHSVYCSTKGALDMLTKMMALELGPHKIRVNAVNPTVVMT
SMGRTNWSDPHKAKVMLDRIPLGKFAEVENVVDAILFLLSHRSNMTTGSTLPVDGGFLVT

>sp|A2QAC0|LAD_ASPNC L-arabinitol 4-dehydrogenase OS=Aspergillus niger (strain CBS 513.88 / FGSC A1513) GN=ladA PE=1 SV=1

MATATVLEKANIGVFTNTKHDLWVADAKPTLEEVKNGQGLQPGEVTIEVRSTGICGSDVHFW
HAGCIGPMIVTGDHILGHESAGQVVAVAPDVTSLKPGDRVAVEPNIICNACEPCLTGRYNGC
ENVQFLSTPPVDGLLRRYVNHPAIWCHKIGDMSYEDGALLEPLSVSLAGIERSGLRLGDPCL
VTGAGPIGLITLLSARAAGASPIVITDIDEGRLEFAKSLVPDVRTYKVQIGLSAEQNAEGIINVFN

DGQGSGPGALRPRIAMECTGVESSVASAIWSVKFGGKVFVIGVGKNEMTVPFMRLSTWEID
LQYQYRYCNTWPRAIRLVRNGVIDLKKLVTHRFLLEDAIKAFETAANPKTGAIKVQIMSSEDDV
KAASAGQKI

>sp|A0QXD8|ELTD_MYCS2 Erythritol/L-threitol dehydrogenase OS=Mycobacterium smegmatis (strain ATCC 700084 / mc(2)155) GN=eltD PE=1 SV=1

MSNQVPEKMQAVVCHGPHDYRLEEVAVPQRKPGEALIRVEAVGICASDLKCYHGAAKFWG
DENRPAWAETMVIPGHEFVGRVVELDDEAAQRWGIAVGDRVVSEQIVPCWECLFCKRGQY
HMCQPHDLYGFKRRTPGAMASYMVYPAEALVHKVSPDIPAQHAAFAEPLSCSLHAVERAQI
TFEDTVVVAGCGPIGLGMIAGAKAKSPMRVIALDMAPDKLKLAEKCGADLTINIAEQDAEKIIK
DLTGGYGADVYIEGTGHTSAVPQGLNLLRKLGRYVEYGVFGSDVTVDWSIISDDKELDVLGA
HLGPYCWPAAIKMIESGALPMDEICTHQFPLTEFQKGLDLVASGKESVKVSLIPA

>sp|O93715|GLCDH_SULSF Glucose 1-dehydrogenase OS=Sulfolobus solfataricus GN=gdh PE=1 SV=1

MKAIIVKPPNAGVQVKDVDEKKLDSYGKIKIRTIYNGICGTDREIVNGKLTLSTLPKGKDFLVLG
HEAIGVVEESYHGFSQGDLVMPVNRRGCGICRNCLVGRPDFCETGEFGEAGIHKMDGFMR
EWWYDDPKYLVKIPKSIEDIGILAQPLADIEKSIEEILEVQKRVPVWTCDDGTLNCRKVLVVGT
GPIGVLFTLLFRTYGLEVWMANRREPTEVEQTVIEETKTNYYNSSNGYDKLKDSVGKFDVIID
ATGADVNILGNVIPLLGRNGVLGLFGFSTSGSVPLDYKTLQEIVHTNKTIIGLVNGQKPHFQQA
VVHLASWKTLYPKAAKMLITKTVSINDEKELLKVLREKEHGEIKIRILWE

>sp|Q96V44|LAD_HYPJE L-arabinitol 4-dehydrogenase OS=Hypocrea jecorina GN=lad1 PE=1 SV=1

MSPSAVDDAPKATGAAISVKPNIGVFTNPKHDLWISEAEPSADAVKSGADLKPGEVTIAVRST
GICGSDVHFWHAGCIGPMIVEGDHILGHESAGEVIAVHPTVSSLQIGDRVAIEPNIICNACEPC
LTGRYNGCEKVEFLSTPPVPGLLRRYVNHPAVWCHKIGNMSWENGALLEPLSVALAGMQR
AKVQLGDPVLVCGAGPIGLVSMLCAAAAGACPLVITDISESRLAFAKEICPRVTTHRIEIGKSA
EETAKSIVSSFGGVEPAVTLECTGVESSIAAAIWASKFGGKVFVIGVGKNEISIPFMRASVREV
DIQLQYRYSNTWPRAIRLIESGVIDLSKFVTHRFPLEDAVKAFETSADPKSGAIKVMIQSLD

>sp|G0RH19|LXR3_HYPJQ L-xylulose reductase OS=Hypocrea jecorina (strain QM6a) GN=lxr3 PE=1 SV=1

MKNGAFPHDNAAVPNVERVLPLFSLKGRTAIVSGAGAGIGLAVAQAFAEAGANVAIWYNSNK
QAVTSAEDIAKTYGVKCKAYQVNVTSAEAVDKAITEIIKEFNGRLDVFVANSGITWTEGAFIDG
SVESARNVMSVNVDGVMWCAKSAGAHFRRQKEEGTTIDGKPLDNFIAGSFIATASMSGSIV
NVPQLQAVYNSSKAAVIHFCKSLAVEWTGFARVNTVSPGYIITEISNFVPPETKTLWKDKIVM
GREGRVGELKGAYLYLASDASSYTTGLDMIVDGGYSLP

>sp|Q876L8|XYL1_HYPJE NAD(P)H-dependent D-xylose reductase xyl1 OS=Hypocrea jecorina GN=xyl1 PE=1 SV=1

MASPTLKLNSGYDMPQVGFGLWKVDNAVCADTVYNAIKAGYRLFDGACDYGNEKECGEGV
ARAIKDGLVKREDLFIVSKLWQTFHDEDKVEPITRRQLADWQIDYFDLFLVHFPAALEYVDPS

VRYPPGWFYDGKSEVRWSKTTTLQQTWGAMERLVDKGLARSIGVSNYQAQSVYDALIYARI
KPATLQIEHHPYLQQPDLVSLAQTEGIVVTAYSSFGPTGFMELDMPRAKSVAPLMDSPVIKAL
ADKHRRTPAQVLLRWATQRGIAVIPKTSRPEVMAQNLDNTSFDLDSEDLAKIADMDLNIRFNK
PTNYFSANKLYLFG

>sp|P37680|SGBE_ECOLI L-ribulose-5-phosphate 4-epimerase SgbE OS=Escherichia coli (strain K12) GN=sgbE PE=1 SV=1

MLEQLKADVLAANLALPAHHLVTFTWGNVSAVDETRQWMVIKPSGVEYDVMTADDMVVVEI
ASGKVVEGSKKPSSDTPTHLALYRRYAEIGGIVHTHSRHATIWSQAGLDLPAWGTTHADYFY
GAIPCTRQMTAEEINGEYEYQTGEVIIETFEERGRSPAQIPAVLVHSHGPFAWGKNAADAVH
NAVVLEECAYMGLFSRQLAPQLPAMQNELLDKHYLRKHGANAYYGQ

>sp|P39306|ULAF_ECOLI L-ribulose-5-phosphate 4-epimerase UlaF OS=Escherichia coli (strain K12) GN=ulaF PE=1 SV=1

MQKLKQQVFEANMELPRYGLVTFTWGNVSAIDRERGLVVIKPSGVAYETMKAADMVVVDMS
GKVVEGEYRPSSDTATHLELYRRYPSLGGIVHTHSTHATAWAQAGLAIPALGTTHADYFFGD
IPCTRGLSEEEVQGEYELNTGKVIIETLGNAEPLHTPGIVVYQHGPFAWGKDAHDAVHNAVV
MEEVAKMAWIARGINPQLNHIDSFLMNKHFMRKHGPNAYYGQK

>sp|P0AFP4|YBBO_ECOLI Uncharacterized oxidoreductase YbbO OS=Escherichia coli (strain K12) GN=ybbO PE=3 SV=1

MTHKATEILTGKVMQKSVLITGCSSGIGLESALELKRQGFHVLAGCRKPDDVERMNSMGFTG
VLIDLDSPESVDRAADEVIALTDNCLYGIFNNAGFGMYGPLSTISRAQMEQQFSANFFGAHQL
TMRLLPAMLPHGEGRIVMTSSVMGLISTPGRGAYAASKYALEAWSDALRMELRHSGIKVSLI
EPGPIRTRFTDNVNQTQSDKPVENPGIAARFTLGPEAVVDKVRHAFISEKPKMRYPVTLVTW
AVMVLKRLLPGRVMDKILQG

>tr|A2QBD7|A2QBD7_ASPNC Aspergillus niger contig An01c0480, genomic contig OS=Aspergillus niger (strain CBS 513.88 / FGSC A1513) GN=An01g14880 PE=4 SV=1

MSLPSHFTINTGAKIPAVGFGTWQAKPLEVENAVEVALREGYRHIDCAAIYRNETEVGNGIRK
SGVPREEIFITGKLWNTKHAPEDVEPALDKTLQDLGVAYLDLYLMHWPCAFKGGDKWFPLN
DDGVFDLANIDYITTYRAMEKLLATGKVRAIGVSNFNIRRLEELLGQVSVVPAVNQIEAHPYLQ
QPDLLQFCQSKGILIEAYSPLGNNQTGEPRTVDDPLVHRVAGELSLDPGPLLASWGVQRGT
VVLSKSVTPARIAANLRVRALPEGAFAQLNSLERHKRFNFPGHWGYDIFEEVGEEAVRQTAL
AAGPSNKVKFTV

>tr|Q9I1Q0|Q9I1Q0_PSEAE Probable aldehyde dehydrogenase OS=Pseudomonas aeruginosa (strain ATCC 15692 / DSM 22644 / CIP 104116 / JCM 14847 / LMG 12228 / 1C / PRS 101 / PAO1) GN=PA2217 PE=4 SV=1

MTAILGHNFIGGARSAAGTLFLQSLDAASGEALPYRFVQATPEEVDAAAEAAASAYPHYRQL
PASRRAEFLDTIAAELDALDDDFVAIVCRETALPATRIQGERSRTSGQLRLFAEVLRRGDFHG
ARIDRARPERKPLPRVDLRQCRIGLGPVAVFGASNFPLAFSTAGGDTAAALAAGCPVVFKAH
SGHMATAERVAAAILRAAERTGMPAGVFNMIYGGGVGERLVRHPAIQAVGFTGSLKGGRAL

CDMAAARAQPIPVFAEMSSINPVVLLPAALKKRGEAVADELSASVVLGCGQFCTNPGLVIGIR
SAQFSAFLERFAARMDDQPAQTMLNTGTLASYEKGLAALHAHPRVRHLAGQPQEGRQARP
QLFQADVSLLLEGDELLQEEVFGPASVVVEVADHAELKRALHGLHGQLTATLIAEAEDLASFA
DLVPLLEQKAGRLLLNGYPTGVEVCDAMVHGGPYPATSDARGTSVGTLAIDRFLRPVCYQN
YPDAWLPEALKDGNPLGIARLVDGIVTRAAVA

>tr|Q9I1Q1|Q9I1Q1_PSEAE Uncharacterized protein OS=Pseudomonas aeruginosa (strain ATCC 15692 / DSM 22644 / CIP 104116 / JCM 14847 / LMG 12228 / 1C / PRS 101 / PAO1) GN=PA2216 PE=4 SV=1

MRLIQFEDRAGQRRVGVVEGAGIQVLRGVRSTRELGLAAIRAGSGLHDEVLRRGSEPGPDY
AGLLEEGRVLPPLDHDDPAHCLVSGTGLTHLGSAATRDRMHQQNQGDETALTDTMRIFRW
GLEGGKPPAGQVGAQPEWFYKGDGGIVVRPGADFPLPAFAEDAGEEPELVGLYLIGDDRRP
YRLGYALGNEFSDHLMERRNYLYLAHSKLRACCYGPELRVGELPRHLQGESRILRDGEVLW
QQAFLSGEDNMCHSLENLEYHHFKYAQFLRPGDVHVHYFGTATLSFADGIKAAPGDVFEIAM
AEFGAPLRNGIAAVETALTPGRVVAL

>tr|Q97VG1|Q97VG1_SULSO Muconate cycloisomerase related protein OS=Sulfolobus solfataricus (strain ATCC 35092 / DSM 1617 / JCM 11322 / P2) GN=SSO2665 PE=4 SV=1

MTKISEIEAYILGKEVTSAQWASLMVLVRVTTNDGRVGWGETVSALRAEAVANFVKKINTVLK
GNDVFNVEKNRLEWYKHDFNMTISLESTTAYSAVDIASWDIIGKELGAPLYKLLGGKTRDKVL
VYANGWYQNCVKPEDFAEKAKEIVKMGYKALKFDPFGPYFNDISKKGLDIAEERVKAVREAV
GDNVDILIEHHGRFNANSAIMIAKRLEKYNPLFMEEPIHPEDVEGLRKYRNNTSLRIALGERIIN
KQQALYFMKEGLVDFLQADLYRIGGVTETKKVVGIAETFDVQMAFHNAQGPILNAVTLQFDA
FIPNFLIQESFYDWFPSWKRELIYNGTPIDNGYAIIPERPGLGVEVNEKMLDSLKVKGEEYFNP
EEPVWVVKGTWRDY

>sp|Q8X167|XKS1_ASPNG D-xylulose kinase A OS=Aspergillus niger GN=xkiA PE=1 SV=1

MQGPLYIGFDLSTQQLKGLVVNSDLKVVYVSKFDFDADSRGFPIKKGVLTNEAEHEVFAPVA
LWLQALDGVLEGLRKQGMDFSQIKGISGAGQQHGSVYWGENAEKLLKELDASKTLEEQLDG
AFSHPFSPNWQDSSTQKECDEFDAALGGQSELAFATGSKAHHRFTGPQIMRFQRKYPDVY
KKTSRISLVSSFIASLFLGHIAPMDISDVCGMNLWNIKKGAYDEKLLQLCAGSSGVDDLKRKL
GDVPEDGGIHLGPIDRYYVERYGFSPDCTIIPATGDNPATILALPLRASDAMVSLGTSTTFLMS
TPSYKPDPATHFFNHPTTAGLYMFMLCYKNGGLARELVRDAVNEKLGEKPSTSWANFDKVT
LETPPMGQKADSDPMKLGLFFPRPEIVPNLRSGQWRFDYNPKDGSLQPSNGGWDEPFDEA
RAIVESQMLSLRLRSRGLTQSPGEGIPAQPRRVYLVGGGSKNKAIAKVAGEILGGSEGVYKL
EIGDNACALGAAYKAVWAMERAEGQTFEDLIGKRWHEEEFIEKIADGYQPGVFERYGQAVE
GFEKMELEVLRQEGKH

>sp|A2Q8B5|XYL1_ASPNC Probable NAD(P)H-dependent D-xylose reductase xyl1 OS=Aspergillus niger (strain CBS 513.88 / FGSC A1513) GN=xyl1 PE=3 SV=1

MASPTVKLNSGYDMPLVGFGLWKVNNDTCADQIYHAIKEGYRLFDGACDYGNEVEAGQGIA
RAIKDGLVKREELFIVSKLWNSFHDGDRVEPICRKQLADWGIDYFDLYIVHFPISLKYVDPAVR
YPPGWKSEKDELEFGNATIQETWTAMESLVDKKLARSIGISNFSAQLVMDLLRYARIRPATLQ

IEHHPYLTQTRLVEYAQKEGLTVTAYSSFGPLSFLELSVQNAVDSPPLFEHQLVKSIAEKHGR
TPAQVLLRWATQRGIAVIPKSNNPQRLKQNLDVTGWNLEEEEIKAISGLDRGLRFNDPLGYG
LYAPIF

>sp|Q876L8|XYL1_HYPJE NAD(P)H-dependent D-xylose reductase xyl1 OS=Hypocrea jecorina GN=xyl1 PE=1 SV=1

MASPTLKLNSGYDMPQVGFGLWKVDNAVCADTVYNAIKAGYRLFDGACDYGNEKECGEGV
ARAIKDGLVKREDLFIVSKLWQTFHDEDKVEPITRRQLADWQIDYFDLFLVHFPAALEYVDPS
VRYPPGWFYDGKSEVRWSKTTTLQQTWGAMERLVDKGLARSIGVSNYQAQSVYDALIYARI
KPATLQIEHHPYLQQPDLVSLAQTEGIVVTAYSSFGPTGFMELDMPRAKSVAPLMDSPVIKAL
ADKHRRTPAQVLLRWATQRGIAVIPKTSRPEVMAQNLDNTSFDLDSEDLAKIADMDLNIRFNK
PTNYFSANKLYLFG

>sp|P09099|XYLB_ECOLI Xylulose kinase OS=Escherichia coli (strain K12) GN=xylB PE=1 SV=1

MYIGIDLGTSGVKVILLNEQGEVVAAQTEKLTVSRPHPLWSEQDPEQWWQATDRAMKALGD
QHSLQDVKALGIAGQMHGATLLDAQQRVLRPAILWNDGRCAQECTLLEARVPQSRVITGNL
MMPGFTAPKLLWVQRHEPEIFRQIDKVLLPKDYLRLRMTGEFASDMSDAAGTMWLDVAKRD
WSDVMLQACDLSRDQMPALYEGSEITGALLPEVAKAWGMATVPVVAGGGDNAAGAVGVG
MVDANQAMLSLGTSGVYFAVSEGFLSKPESAVHSFCHALPQRWHLMSVMLSAASCLDWAA
KLTGLSNVPALIAAAQQADESAEPVWFLPYLSGERTPHNNPQAKGVFFGLTHQHGPNELAR
AVLEGVGYALADGMDVVHACGIKPQSVTLIGGGARSEYWRQMLADISGQQLDYRTGGDVG
PALGAARLAQIAANPEKSLIELLPQLPLEQSHLPDAQRYAAYQPRRETFRRLYQQLLPLMA

>sp|P00944|XYLA_ECOLI Xylose isomerase OS=Escherichia coli (strain K12) GN=xylA PE=3 SV=1

MQAYFDQLDRVRYEGSKSSNPLAFRHYNPDELVLGKRMEEHLRFAACYWHTFCWNGADM
FGVGAFNRPWQQPGEALALAKRKADVAFEFFHKLHVPFYCFHDVDVSPEGASLKEYINNFA
QMVDVLAGKQEESGVKLLWGTANCFTNPRYGAGAATNPDPEVFSWAATQVVTAMEATHKL
GGENYVLWGGREGYETLLNTDLRQEREQLGRFMQMVVEHKHKIGFQGTLLIEPKPQEPTKH
QYDYDAATVYGFLKQFGLEKEIKLNIEANHATLAGHSFHHEIATAIALGLFGSVDANRGDAQL
GWDTDQFPNSVEENALVMYEILKAGGFTTGGLNFDAKVRRQSTDKYDLFYGHIGAMDTMAL
ALKIAARMIEDGELDKRIAQRYSGWNSELGQQILKGQMSLADLAKYAQEHHLSPVHQSGRQ
EQLENLVNHYLFDK

>sp|Q3SYZ6|XYLB_BOVIN Xylulose kinase OS=Bos taurus GN=XYLB PE=2 SV=1

MAERAARHCCLGWDFSTQQVKVVAVDAELSVFYEDSVHFDRDLVEFGTQGGVHVHKDGLT
VTSPVLMWVQALDIILEKMKASGFDFSQVLALSGAGQQHGSVYWKTGASQVLTSLSPDLPL
REQLQACFSISNCPVWMDSSTAAQCRQLEAAVGGAQALSLLTGSRAYERFTGNQIAKIYQQ
NPEAYSHTERISLVSSFAASLFLGSYSPVDYSDGSGMNLLQIQDKVWSQACLGACAPRLEEK
LGRPVPSCSIVGAISSYFVQRYGFPPECKVVAFTGDNPASLAGMRLEEGDIAVSLGTSDTLFL
WLQEPTPALEGHIFCNPVDPQHYMALLCFKNGSLMREKIRDESASGSWSKFSKALQSTGMG

NSGNLGFYFDVMEITPEIIGRHRFTAENHEVSAFPQDVEIRALIEGQFMAKKIHAEALGYRVM
PKTKILATGGASHNRDILQVLADVFGAPVYVIDTANSACVGSAYRAFHGPSLLCLVSIY

>sp|P0AFP4|YBBO_ECOLI Uncharacterized oxidoreductase YbbO OS=Escherichia coli (strain K12) GN=ybbO PE=3 SV=1

MTHKATEILTGKVMQKSVLITGCSSGIGLESALELKRQGFHVLAGCRKPDDVERMNSMGFTG
VLIDLDSPESVDRAADEVIALTDNCLYGIFNNAGFGMYGPLSTISRAQMEQQFSANFFGAHQL
TMRLLPAMLPHGEGRIVMTSSVMGLISTPGRGAYAASKYALEAWSDALRMELRHSGIKVSLI
EPGPIRTRFTDNVNQTQSDKPVENPGIAARFTLGPEAVVDKVRHAFISEKPKMRYPVTLVTW
AVMVLKRLLPGRVMDKILQG

>Ss-LADH_tr|A3LNE3|A3LNE3_PICST Aldehyde dehydrogenase OS=Scheffersomyces stipitis (strain ATCC 58785 / CBS 6054 / NBRC 10063 / NRRL Y-11545) OX=322104 GN=ALD5 PE=3 SV=1

MSLPLFVPIKLPNGTTYEQPTGLFINNEFVQSKSKKTFGTVSPSTEEEITQVYEAFSEDIDDAV
EAATAAFHSSWSTSDPQVRMKVLYKLADLIDEHADTLAHIEALDNGKSLMCSKGDVALTAAY
FRSCAGWTDKIKGSVIETGDTHFNYTRREPIGVCGQIIPWNFPLLMASWKLGPVLCTGCTTVL
KTAESTPLSALYLASLIKEAGAPPGVVNVVSGFGPTAGAPISSHPKIKKVAFTGSTATGRHIMK
AAAESNLKKVTLELGGKSPNIVFDDADVKSTIQHLVTGIFYNTGEVCCAGSRIYVQEGIYDKIV
SEFKNAAESLKIGDPFKEDTFMGAQTSQLQLDKILKYIDIGKKEGATVITGGERFGNKGYFIKP
TIFGDVKEDHQIVRDEIFGPVVTITKFKTVEEVIALANDSEYGLAAGVHTTNLSTAISVSNKINS
GTIWVNTYNDFHPMVPFGGYSQSGIGREMGEEALDNYTQVKAVRIGLSQ

>Ss-LRA1_sp|A3LZU7|RM1DH_PICST L-rhamnose-1-dehydrogenase OS=Scheffersomyces stipitis (strain ATCC 58785 / CBS 6054 / NBRC 10063 / NRRL Y-11545) OX=322104 GN=DHG2 PE=1 SV=2

MTGLLNGKVVAITGGVTGIGRAIAIEMARNGAKVVVNHLPSEEQAQLAKELKEEISDGENNVL
TIPGDISLPETGRRIVELAVEKFGEINVFVSNAGVCGFREFLEITPETLFQTVNINLNGAFFAIQA
AAQQMVKQGKGGSIIGISSISALVGGAHQTHYTPTKAGILSLMQSTACALGKYGIRCNAILPGT
ISTALNEEDLKDPEKRKYMEGRIPLGRVGDPKDIAGPAIFLASDMSNYVNGAQLLVDGGLFVN
LQ

>Dh-LRA2_tr|Q6BQZ8|Q6BQZ8_DEBHA DEHA2E01078p OS=Debaryomyces hansenii (strain ATCC 36239 / CBS 767 / JCM 1990 / NBRC 0083 / IGC 2968) OX=284592 GN=DEHA2E01078g PE=4 SV=2

MPSKKYKIIDSHVHLFAKRNFKLLKFDEAHPLHSDFRLDEYLKYSMNEEFQIDGLVFIETDPIA
DLSKALEGCEYPIQEYLYVARNITGNLLPDEGETSELKQNFIKAIVPWAPMPLGKSSVSSYVE
MLKSRSSTDEFNLVKGFRYLVQDKLPNTMLQRDFVESLKWLDDNNFIFDWGIDLRCGGLWQ
FEETIEVLKQVPNLKYVINHLTKPNLSIDPTKIEENDEFLQWKNYMKQIFVNSPNSYMKLSGGF
SELPSEIIENRDKCAEYIYPWFKVCFDLWNVDRTIWASNWPVCTLTAGEDLTSKWFEVTEML
FDKIELNEESRKKIYGTNYLKAYNLI

>Ss-LRA4_tr|A3LZU9|A3LZU9_PICST L-KDR aldolase OS=Scheffersomyces stipitis (strain ATCC 58785 / CBS 6054 / NBRC 10063 / NRRL Y-11545) OX=322104 GN=PICST_64442 PE=3 SV=1

MTISAALPKRGVYTPVPTFFKKDLHTIDYDSQIEHAKFLQQNGITGLVLLGSTGENSHLTRKER
IELVSTIHEELPDFPLMAGVAQNSVEDAIEEILQLKNAGAQHALVLPSSYFGASIKQQGIIDWYT
EVADNASLPVLIYVYPGVSNNISIDPRTIKKLSAHPNIVGAKISHGDVSHHAIIGLDQEIAANQFI
TLTGLGQILLPVLVVGIQGTVDALCGAFPKIYVKLLENYDKGDLRAAAELQLVISRAEELVVKF
GVVGIKKAIHFATGIGETYLGRAPLTQDVNDADWKSYNDYLLGIVSVESTL

>sp|Q38707|MTDH_APIGR Mannitol dehydrogenase OS=Apium graveolens OX=4045 GN=MTD PE=1 SV=1

MAKSSEIEHPVKAFGWAARDTTGLLSPFKFSRRATGEKDVRLKVLFCGVCHSDHHMIHNNW
GFTTYPIVPGHEIVGVVTEVGSKVEKVKVGDNVGIGCLVGSCRSCESCCDNRESHCENTIDT
YGSIYFDGTMTHGGYSDTMVADEHFILRWPKNLPLDSGAPLLCAGITTYSPLKYYGLDKPGT
KIGVVGLGGLGHVAVKMAKAFGAQVTVIDISESKRKEALEKLGADSFLLNSDQEQMKGARSS
LDGIIDTVPVNHPLAPLFDLLKPNGKLVMVGAPEKPFELPVFSLLKGRKLLGGTINGGIKETQE
MLDFAAKHNITADVEVIPMDYVNTAMERLVKSDVRYRFVIDIANTMRTEESLGA

>Ss_agaK_sp|A0KYQ6|AGAK_SHESA N-acetylgalactosamine kinase AgaK OS=Shewanella sp. (strain ANA-3) OX=94122 GN=agaK PE=1 SV=1

MYYGLDIGGTKIELAIFDTQLALQDKWRLSTPGQDYSAFMATLAEQIEKADQQCGERGTVGIA
LPGVVKADGTVISSNVPCLNQRRVAHDLAQLLNRTVAIGNDCRCFALSEAVLGVGRGYSRVL
GMILGTGTGGGLCIDGKLYLGANRLAGEFGHQGVSANVACRHQLPLYVCGCGLEGCAETYV
SGTGLGRLYQDIAGQTADTFAWLNALRCNDPLAIKTFDTYMDILGSLMASLVLAMDPDIIVLG
GGLSEVEEILAALPQATKAHLFDGVTLPQFKLADFGSASGVRGAALLGHGLDAGISYEA

>Ss_agaA_sp|A0KYQ5|AGAA2_SHESA N-acetylgalactosamine-6-phosphate deacetylase OS=Shewanella sp. (strain ANA-3) OX=94122 GN=agaAII PE=1 SV=1

MKPNTDFMLIADGAKVLTQGNLTEHCAIEVSDGIICGLKSTISAEWTADKPHYRLTSGTLVAG
FIDTQVNGGGGLMFNHVPTLETLRLMMQAHRQFGTTAMLPTVITDDIEVMQAAADAVAEAID
CQVPGIIGIHFEGPHLSVAKRGCHPPAHLRGITEREWLLYLRQDLGVRLITLAPESVTPEQIKR
LVASGAIISLGHSNADGETVLKAIEAGASGFTHLYNGMSALTSREPGMVGAAFASENTYCGIIL
DGQHVHPISALAAWRAKGTEHLMLVTDAMSPLGSDQTEFQFFDGKVVREGMTLRDQHGSL
AGSVLDMASAVRYAATELNLGLSNAVQMATRTPAEFIQRPQLGDIAEGKQADWVWLDDDQ
RVLAVWIAGELLYQAEQARFA

>Ss_agaS_sp|A0KYQ7|AGAS_SHESA D-galactosamine-6-phosphate deaminase AgaS OS=Shewanella sp. (strain ANA-3) OX=94122 GN=agaS PE=1 SV=1

MLTSPLSPFEHEDSNLLLSAEQLTQYGAFWTAKEISQQPKMWRKVSEQHSDNRTIAAWLTPI
LAKPQLRIILTGAGTSAYIGDVLAAHIQQHLPLATQQVEAISTTDIVSHPELYLRGNIPTLLISYG
RSGNSPESMAAVELAEQLVDDCYHLAITCNGQGKLANYCADKSHCYLYKLPDETHDVSFAM
TSSFTCMYLATLLIFAPNSQALMQCIEMAEHILTERLADIRLQSEQPSKRVVFLGGGPLKAIAQ

EAALKYLELTAGQVVSAFESPLGFRHGPKSLVDSHTQVLVMMSSDPYTRQYDNDLIQELKRD
NQALSVLTLSEELLTGSSGLNEVWLGLPFILWCQILAIYKAIQLKVSPDNPCPTGQVNRVVQG
VNVYPFVK

>AAA93234.2 amygdalin hydrolase isoform AH I precursor [Prunus serotina]

MATKLGSLLLCALLLAGFALTNSKAAKTDPPIHCASLNRSSFDALEPGFIFGTASAAYQFEGA
AKEDGRGPSIWDTYTHNHSERIKDGSNGDVAVDQYHRYKEDVRIMKKMGFDAYRFSISWSR
VLPNGKISGGVNEDGIKFYNNLINEILRNGLKPFVTIYHWDLPQALEDEYGGFLSPNIVDHFRD
YANLCFKKFGDRVKHWITLNEPYTFSSSGYAYGVHAPGRCSAWQKLNCTGGNSATEPYLVT
HHQLLAHAAAVKLYKDEYQASQNGLIGITLVSPWFEPASEAEEDINAAFRSLDFIFGWFMDPL
TNGNYPHLMRSIVGERLPNFTEEQSKLLKGSFDFIGLNYYTTRYASNAPKITSVHASYITDPQ
VNATAELKGVPIGPMAASGWLYVYPKGIHDLVLYTKEKYNDPLIYITENGVDEFNDPKLSMEE
ALKDTNRIDFYYRHLCYLQAAIKKGSKVKGYFAWSFLDNFEWDAGYTVRFGINYVDYNDNLK
RHSKLSTYWFTSFLKKYERSTKEIQMFVESKLEHQKFESQMMNKVQSSLAVVV

>ACO22019.1 beta-glucosidase (Gentiobiase) (Cellobiase) (Beta-D-glucoside glucohydrolase) (Amygdalase) [Streptococcus pneumoniae P1031]

MTIFPDDFLWGGAVAANQVEGAYNEDGKGLSVQDVLPKGGLGEATENPTEDNLKLIGIDFYH
KYKEDISLFSEMGFNVFRTSIAWSRIFPKGDEEEPNEAGLKYYDELFDELHAHGIEPLVTLSH
YETPLYLARKYHGWIDRRMIHFYEKFARTVLERYKDKVKYWLTFNEVNSVLELPFTSGGIDIP
KENLSKQELYQAIHHELVASSLVTKIAREINSEFKVGCMVLAMPAYPMTPNPKDVWATHEYE
NLNYLFSDVHVRGYYPNYAKRYFKENDINIEFAAEDAELLKNYTVDFLSFSYYMSVTQSALPT
QYNSGEGNIIGGLVNPYLESSEWGWQIDPIGLRIILNRYYDRYQIPLFIVENGLGAKDQLIKDEF
NNLTVQDDYRIQYMKEHLLQVAEALQDGVEIMGYTSWGCIDCVSMSTAQLSKRYGLIYVDRN
DDGNGTFNRYKKMSFTWYKGVIESNGESLFK

>ACK40981.1 beta-glucosidase (Gentiobiase) (Cellobiase) (Beta-D-glucoside glucohydrolase) (Amygdalase) [Listeria monocytogenes HCC23]

MTESKFPKGFLWGGAVAANQCEGAYLEDGKGLSLVDILPTVEDGRWEALFNPSKALATDYG
FYPSHESIDFYHRYKEDIKLMAEMGFKCFRMSISWPRIFPNGDETTPNEKGLAFYDAVFDEC
HKYGIEPVVTINHFDTPLEVFKKYGGWKNRKCIDFYLNFCEAIFTRYKDKVKYWMTFNEINMIL
HLPYIGGGLDVTKEANPEEVKYQAAHHQLVASALATKLGHEINPENQIGCMLAAGNTYPMTC
NPKDVWKSIEADREGYFFIDVQARGYYPSYTKRFFKEHNINIKMEDGDLDALRDHTVDYVAF
SYYSSRLTSADPEKNKETEGNVFATLKNPYLKASEWGWQIDPLGLRITMNTIYDRYQKPLFIV
ENGLGAVDTVAEDGSITDDYRIDYMREHVREMGEAIEDGVELLGYTPWGCIDLVSAGSGEM
KKRYGFIYVDRDNKGNGTLNRSKKKSFDWYKKVIETNGKDID

>D-ribose_pyranase sp|P04982|RBSD_ECOLI D-ribose pyranase OS=Escherichia coli (strain K12) OX=83333 GN=rbsD PE=1 SV=3

MKKGTVLNSDISSVISRLGHTDTLVVCDAGLPIPKSTTRIDMALTQGVPSFMQVLGVVTNEMQ
VEAAIIAEEIKHHNPQLHETLLTHLEQLQKHQGNTIEIRYTTHEQFKQQTAESQAVIRSGECSP
YANIILCAGVTF

>ribokinase_S_cerevisiae EDN62156.1 ribokinase [Saccharomyces cerevisiae YJM789]

MGITVIGSLNYDLDTFTDRLPNAGETFRANHFETHAGGKGLNQAAAIGKLKNPSSRYSVRMI
GNVGNDTFGKQLKDTLSDCGVDITHVGTYEGINTGTATILIEEKAGGQNRILIVEGANSKTIYD
SKQLCEIFPEGKEEEEYVVFQHEIPDPLSIIKWIHANRPNFQIVYNPSPFKAMRKKDWELVDLL
VVNEIEGLQIVESVFDNELVEEIREKIKDDFLGEYRKICELLYEKLMNRKKRGIVVMTLGSKGV
LFCSHESPEVQFLPAIENVSVVDTTGAGDTFLGGLVTQLYQGETLSTAIKFSTLASSLTIQRKG
AAESMPLYKDVQKDA

>phosphoribomutase_S_cerevisiae NP_014005.1 phosphoribomutase PRM15
[Saccharomyces cerevisiae S288C]

MLQGILETVPSDLKDPISLWFKQDRNPKTIEEVTALCKKSDWNELHKRFDSRIQFGTAGLRSQ
MQAGFSRMNTLVVIQASQGLATYVRQQFPDNLVAVVGHDHRFHSKEFARATAAAFLLKGFK
VHYLNPDHEFVHTPLVPFAVDKLKASVGVMITASHNPKMDNGYKVYYSNGCQIIPPHDHAIS
DSIDANLEPWANVWDFDDVLNKALKQGKLMYSREEMLKLYLEEVSKNLVEINPLKLEVKAKP
WFVYTPMHGVGFDIFSTIVKKTLCLVEGKDYLCVPEQQNPDPSFPTVGFPNPEEKGALDIGIN
LAEKHDIDLLVANDPDADRFSVAVKDMQSGEWRQLTGNEIGFLFAFYEYQKYKSMDKEFQH
VHPLAMLNSTVSSQMIKKMAEIEGFHYEDTLTGFKWIGNRAILLEKKGYYVPFGFEEAIGYMF
PAMEHDKDGISASIVFLQAYCKWKIDHNLDPLNVLENGFKKYGVFKEYNGYYVVPNPTVTKDI
FDYIRNVYTPEGASYPSSIGEEIEVLYYRDLTTGYQSDTINHKPTLPVDPTSQMITVSARPSNG
SENEHIRFTIRGSGTEPKLKVYIEACANEEQRASFLAKLTWNVLRREWFRPDEMNIVTKF

>Acetaldehyde_dehydrogenase_S_cerevisiae AAB68304.1 Ald6p: Acetaldehyde
dehydrogenase [Saccharomyces cerevisiae]

MTKLHFDTAEPVKITLPNGLTYEQPTGLFINNKFMKAQDGKTYPVEDPSTENTVCEVSSATTE
DVEYAIECADRAFHDTEWATQDPRERGRLLSKLADELESQIDLVSSIEALDNGKTLALARGDV
TIAINCLRDAAAYADKVNGRTINTGDGYMNFTTLEPIGVCGQIIPWNFPIMMLAWKIAPALAMG
NVCILKPAAVTPLNALYFASLCKKVGIPAGVVNIVPGPGRTVGAALTNDPRIRKLAFTGSTEVG
KSVAVDSSESNLKKITLELGGKSAHLVFDDANIKKTLPNLVNGIFKNAGQICSSGSRIYVQEGI
YDELLAAFKAYLETEIKVGNPFDKANFQGAITNRQQFDTIMNYIDIGKKEGAKILTGGEKVGDK
GYFIRPTVFYDVNEDMRIVKEEIFGPVVTVAKFKTLEEGVEMANSSEFGLGSGIETESLSTGL
KVAKMLKAGTVWINTYNDFDSRVPFGGVKQSGYGREMGEEVYHAYTEVKAVRIKL

>deoxyribose-phosphate_aldolase_P_lactucaedebilis ORY78125.1 deoxyribose-phosphate
aldolase [Protomyces lactucaedebilis]

MNRLIDHTILKPDATKAEVEKICDEALALETATVCVNTRWLPLVSKKLANSNVLPIAVVGFPLG
ACLTEAKVFETKLAIQQGAKEIDMVIDVGALKDGEVEHVEKDIHAVVQAAGNIPVKVILETCLLT
DEQKRTACKLCKKAGAAFVKTSTGFSKSGATVADTKLMREEVGKEMGVKASGGIRTFKDAQ
AMVDAGASRIGASASVAIMAEANASSNV

>aldehyde-alcohol_dehydrogenase_Fusarium KLP10254.1 aldehyde-alcohol dehydrogenase
[Fusarium fujikuroi]

MTDTLKPYRMSQLEGGHRPGSDAIFNSHSKDGILSALKEWNSQRILLVHSKALAKNTHVISYL
KEALGDRLTNVKEGVGSHSPYSDVIDIAHRITEHKIDCVISVGSGSYSDACKVARLMSATLPA
GFREEDMENLLDQDKGVTPQDKMKKADGVKLILVPTSLSAGEWNHTASCTNSAGKKQHFSL
QDGGAPDLILMDPWVARTSPEKLWMSSGIRAVDHCVETLCNPECKKYPDVQEWCEEALRD
LAKGLVEYKEGLGRGKEGEDELVHGVSKCQTGSRMALMGFIIYRVNMGASHAIGHQLGSVG
KVMHGITSCIMLPPVLRYTKARNPGAQARIVEIFNEALGWEETDASDCVARLVEVTGLPSTLR
DVGVTNNEQIEQIIDKTMTDVMFSFGKILTRKEVSEIVYSTK
